# The timing of human adaptation from Neanderthal introgression

**DOI:** 10.1101/2020.10.04.325183

**Authors:** Sivan Yair, Kristin M. Lee, Graham Coop

**Affiliations:** Center for Population Biology, University of California, Davis; Department of Evolution and Ecology, University of California, Davis; Department of Biological Sciences, Columbia University

## Abstract

Admixture has the potential to facilitate adaptation by providing alleles that are immediately adaptive in a new environment or by simply increasing the long term reservoir of genetic diversity for future adaptation. A growing number of cases of adaptive introgression are being identified in species across the tree of life, however the timing of selection, and therefore the importance of the different evolutionary roles of admixture, is typically unknown. Here, we investigate the spatio-temporal history of selection favoring Neanderthal-introgressed alleles in modern human populations. Using both ancient and present-day samples of modern humans, we integrate the known demographic history of populations, namely population divergence and migration, with tests for selection. We model how a sweep placed along different branches of an admixture graph acts to modify the variance and covariance in neutral allele frequencies among populations at linked loci. Using a method based on this model of allele frequencies, we study previously identified cases of Neanderthal adaptive introgression. From these, we identify cases in which Neanderthal introgressed alleles were quickly beneficial and other cases in which they persisted at low frequency for some time. For some of the alleles that persisted at low frequency, we show that selection likely independently favored them later on in geographically separated populations. Our work highlights how admixture with ancient hominins has contributed to modern human adaptation and contextualizes observed levels of Neanderthal ancestry in present-day and ancient samples.

## 1 Introduction

Within the last decade, population genomic studies have revealed many examples of hybridization that lead to the introgression of genetic material between diverged populations. While these genetic introductions can be ecologically or developmentally maladaptive, a growing number of studies show that natural selection sometimes favors the spread of introgressed alleles (e.g. Whitney et al., 2006; The Heliconius Genome Consortium, 2012; Jones et al., 2018; Oziolor et al., 2019). Introgressed alleles have been identified as contributing to adaptation to new or changing environments, highlighting introgression as a potentially important source of genetic variation for fitness.

Patterns of archaic introgression in modern humans offer some of the most compelling examples of adaptive introgression (reviewed in Racimo et al., 2015). When a subset of modern humans spread out of Africa, likely in the past hundred thousand years, they encountered a broad range of novel environments, including reduced UV exposure, new pathogen pressures, and colder climates. At roughly the same time they experienced new conditions, modern humans outside of Africa were also interbreeding with Neanderthals, who had been living in and adapting to these Eurasian environments for hundreds of thousands of years (Hublin, 2009). The early-generation hybrids of Neanderthals and modern humans may have had low fitness due to the accumulation of weakly deleterious alleles in Neanderthals, who had a small effective population size. The low fitness of these early hybrids, combined with the greater efficacy of purifying selection in modern humans, led to widespread selection against deleterious Neanderthal alleles and linked Neanderthal variation (Harris and Nielsen, 2016; Juric et al., 2016). As a result, Neanderthal alleles segregate around 1-3% frequency genome-wide in present day modern humans, with a depletion in gene rich, regulatory, and low recombination regions (Sankararaman et al., 2014; Schumer et al., 2018; Steinrücken et al., 2018; Petr et al., 2019; Telis et al., 2020).

While, on average, Neanderthal-derived alleles have been selected against, a few alleles exist at high frequency in present-day non-African populations and reflect putative cases of adaptive introgression (Abi-Rached et al., 2011; Sankararaman et al., 2014; Khrameeva et al., 2014; Vernot and Akey, 2014; Dannemann et al., 2016; Deschamps et al., 2016; Gittelman et al., 2016; Racimo et al., 2017; Jagoda et al., 2018; Quach et al., 2016; Sams et al., 2016; Setter et al., 2020). These alleles have been identified based on the characteristic patterns left behind by adaptive introgression, namely high haplotype similarity between Eurasian modern humans and Neanderthals, and the deep divergence of haplotypes within modern human populations. It appears that most sweeps on Neanderthal introgressed alleles were partial sweeps, with selected alleles only reaching frequencies around 30 to 60%. Thus, dips in genetic diversity due to adaptive Neanderthal introgression are dampened relative to hard, full sweeps on *de novo* mutations.

Some of the selected Neanderthal alleles contribute to traits that may have been under selection during modern human expansion out of Africa, such as immunity, skin pigmentation, and metabolism (Abi-Rached et al., 2011; Sankararaman et al., 2014; Khrameeva et al., 2014; Vernot and Akey, 2014; Dannemann et al., 2016; Gittelman et al., 2016; Racimo et al., 2017; Jagoda et al., 2018; Quach et al., 2016). Since modern humans mated with Neanderthals around the same time they were exposed to new environments, a natural assumption is that Neanderthal variation immediately facilitated modern human adaptation. Alternatively, some of the selected Neanderthal introgressed alleles may have contributed to the reservoir of standing genetic variation that became adaptive much later (Jagoda et al., 2018) as human populations were exposed to and created further novel environments. A recent study found evidence of this phenomenon for a Denisovan haplotype contributing to high altitude adaptation on the Tibetan Plateau (Zhang et al., 2020), and so non-immediate selection on introgressed archaic variation may be more common than previously thought.

Population genetics offers a number of approaches to date the timing of selection on alleles. The first broad category of approaches relies on the hitchhiking signal created in surrounding linked variants as the selected alleles sweep up a chunk of the haplotype on which they arose (or introgressed) (Maynard Smith and Haigh, 1974). Second, ancient DNA now offers an opportunity to assess when these selected haplotypes rose in frequency, and the ancient populations where they first achieved high frequency. Currently, methods that use ancient DNA to investigate the temporal history of selection focus on identifying significant increases in selected allele frequencies over time (e.g. Mathieson et al., 2015; Schraiber et al., 2016; Mathieson and Mathieson, 2018). These time series approaches are powerful because they can provide more direct evidence of selection driving allele frequency change, however they are limited to characterizing cases of selection that began after the oldest sampling time and among sampled populations. For studies of modern human evolution, we are currently restricted to learning about adaptation in mostly European populations and within the last ten thousand years, long after hybridization with Neanderthals.

Here, we leverage the advantages of both temporal sampling and patterns of neutral diversity at linked loci (the hitchhiking effect) to infer the time that Neanderthal introgressed alleles became adaptive. Our investigation of the hitchhiking effect allows us to date selection older than the ancient samples, while ancient populations provide useful reference points closer to the sweep time. We use coalescent theory to describe patterns of sequence similarity around a selected site when selection favors introgressed variation at different times since admixture. Our predictions relate sequences among ancient and present day populations by incorporating information about their history of divergence and migration. In addition, we utilize information from partial sweeps by modeling this hitchhiking effect among both selected and non-selected haplotypes. Our approach builds on the model-based inference framework introduced in Lee and Coop (2017), which connects predicted coalescent histories to their corresponding probability distribution of population allele frequencies. Thus, we can distinguish among possible selection times by analyzing allele frequency data from ancient and present day populations.

By providing the age of selection favoring Neanderthal alleles in specific regions, we can determine the context within which Neanderthal alleles facilitated modern human adaptation and in turn narrow their potential phenotypic contributions. If selection was immediate, then the Neanderthal variation was useful in the early Eurasian environments that modern humans experienced, possibly because Neanderthals had been adapting to those same conditions for a long period of time. Otherwise, we have identified Neanderthal haplotypes that were selected during different time periods and in different populations. These latter haplotypes are particularly interesting candidates, as they shed light on the similarities and differences in selection pressures affecting human populations over space and time. Further, they show that admixture between closely related species can provide an important source of standing genetic variation for future adaptation. In this paper, we provide evidence for each of these scenarios in different genomic regions of adaptive introgression.

We begin by introducing the data that we analyzed, followed by an intuitive description of the hitchhiking patterns that we modeled before providing the mathematical details of our model and inference framework. We then show how our method performs on simulations and its results when applied to candidates of adaptive Neanderthal introgression.

## 2 Distribution of adaptive Neanderthal introgressed haplotypes across populations

### Choice of population samples

We applied our method to a set of eight population samples: one archaic (Neanderthals), three present day modern human, and four ancient modern human. For the Neanderthal sample we used the Vindija *33.19* Neanderthal (n=2, for two chromosomes sampled from one individual) because it is high coverage and most closely related to the introgressing population (Prüfer et al., 2017; Mafessoni et al., 2020). Our results only change slightly when the high coverage Altai Neanderthal sample is included in the Neanderthal population, see the section describing the method’s application. The present day populations are from Phase 3 of the 1000 Genomes Project (The 1000 Genomes Project Consortium, 2015): the Yoruba in Ibadan, Nigeria (YRI; n=216), Han Chinese from Beijing (CHB; n=206), and Utah Residents (CEPH) with Northern and Western European Ancestry (CEU; *n* = 198). We chose ancient populations based on previous work about their relationships with present day populations. We included the West Eurasian Upper Paleolithic (EurUP; *n* = 12), who are the oldest samples and basally related to all West Eurasians (mean sampling time *t* = 34kya), and three populations known to be ancestral to present day Europeans: the Mesolithic hunter gatherers from western Europe (WHG; *n* = 108; *t* = 9kya), the Neolithic Anatolian Farmers (EF; *n* = 286; *t* = 7kya), and the Bronze Age Steppe individuals (Steppe; *n* = 38; *t* = 5kya). Each population ancestral to present day Europeans was composed of samples with ≥ 90% inferred ancestry corresponding to that population. Inferred ancestry proportions are from Mathieson et al. (2018)‘s supervised ADMIXTURE analysis using four clusters with fixed membership corresponding to each of these three ancient populations and Eastern Hunter Gatherers. See Table S1 for detailed information on the ancient samples we analyze. One reason for taking this set of ancient populations is that our method requires a known population history, namely the approximate timing and admixture proportions among populations, as well as the timing of divergence among Neanderthal-admixed populations. The primary admixture graph that we used is shown in Figure 1 (Skoglund et al., 2012; Lazaridis et al., 2014; Allentoft et al., 2015; Haak et al., 2015; Mathieson et al., 2015; Fu et al., 2016; Lipson et al., 2017; Sikora et al., 2017; Mathieson et al., 2018). We acknowledge that we take a coarse approach by simplifying human demographic history to a set of ancestral populations, who themselves are products of mixtures of the past.

**Figure 1:**
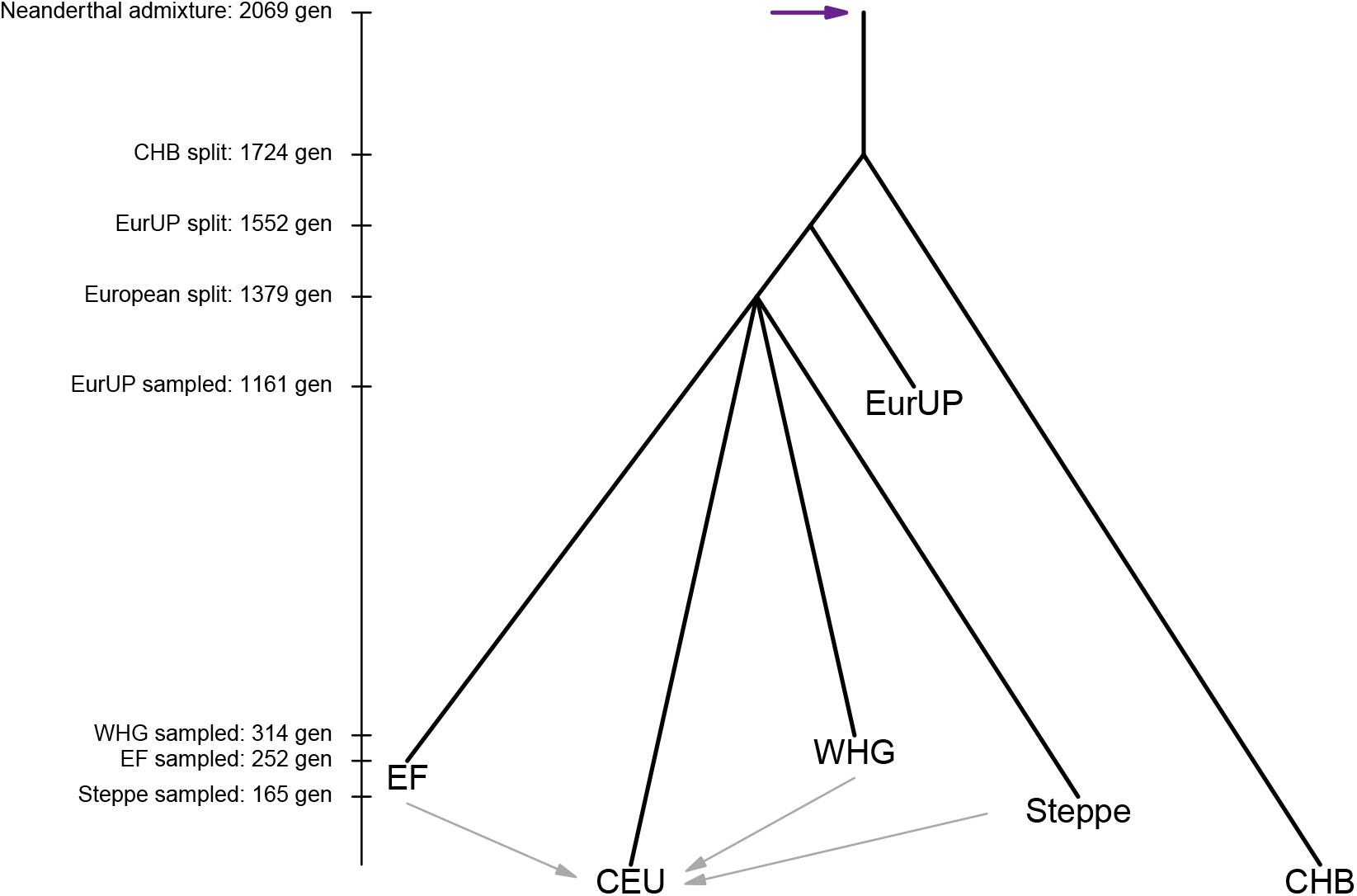
The primary admixture graph that we use in our method. Our method requires the divergence times among admixed modern human populations, the time and proportion of admixture with Neanderthals, and the sampling time of each population. It also requires the timing and proportions of admixture among modern human populations, which we describe in detail in Table S2. We converted many of these estimates from years to generations assuming a generation time of 29 years (Fenner, 2005). Our method is robust to modifications to this admixture graph, which we discuss later.

All ancient modern human samples were typed on a 1240k capture array. Since we needed dense sampling of the neutral loci surrounding the selected site, we imputed genotypes in the ancient modern human samples using Beagle 4.1 (Browning and Browning, 2007, 2016). To ensure imputation accuracy, we selected ancient samples for analysis if they had ≥ 1x coverage. This cutoff was chosen because at this point an imputation procedure very similar to ours can correctly recover around 80-95% of an ancient sample’s heterozygous sites and preserves properties of the genotypic data such as samples’ PCA locations relative to no imputation (Mathieson, 2016). To perform the imputation we followed a procedure similar to that of Mathieson et al. (2015). At the 1240k sites, we computed genotype likelihoods from read counts according to binomial likelihoods of these counts and a small amount of sequencing error. We then ran Beagle twice, first to impute genotypes at just the 1240k sites using the `gl` and `impute=false` arguments, and then to impute the remaining sites using the `gt` and `impute=true arguments`. After each round of imputation we removed polymorphic sites with allelic *r*^2^ < 0.8 (a measure of imputation accuracy). These intermediate filtering steps can improve the imputation accuracy of even ultra-low coverage ancient samples (Hui et al., 2020). To account for some of the uncertainty in the imputation procedure, we calculated population allele frequencies by weighting sample genotypes by their posterior probability, rather than using the maximum likelihood genotypes. We imputed each population separately, used the HapMap Project’s genetic map (The International HapMap Consortium, 2007), and used a reference panel consisting of all East Asian and European 1000 Genomes populations (as multi-population panels can improve imputation accuracy in untyped populations; Huang et al., 2009). Finally, to assess the impact of imputation uncertainty on our results, we also ran our method on bootstrapped datasets, which we describe after introducing the method.

### Choice of Neanderthal introgressed regions

We analyzed genomic regions that were previously identified as putative candidates of adaptive introgression by Racimo et al. (2017). We chose regions with signals of adaptive introgression in European populations, as we have more information on their ancestral populations to infer the age of selection. We removed any regions whose introgressed sequences could not be distinguished from Denisovan ancestry (according to Racimo et al., 2017) and any regions on the X chromosome. To ensure we were capturing the full window of potentially selected sites, we extended the 45kb windows identified by Racimo et al. (2017) by 2 × 10^−2^ cM (20kb if the window has a constant per base pair recombination rate of 10^−8^) and collapsed those that were overlapping or directly adjacent. This resulted in 36 distinct regions, listed in Table S3, out of 50 45kb windows with signals of adaptive Neanderthal introgression in Europeans.

### Neanderthal ancestry over time

As a first step to understand the temporal history of selection, we investigated levels of Neanderthal ancestry in ancient and present day admixed modern human populations within each of the previously specified regions. These levels were determined from ancestry informative sites, which we identified as bi-allelic sites in modern humans that have one allele fixed in Neanderthals (combined Vindija and Altai, two high coverage Neanderthals) and at less than 5% frequency in Yoruba. We included the Altai Neanderthal sample here to increase our confidence that we identified true fixed differences between Neanderthal and modern human lineages. In Figure 2, we show average Neanderthal allele frequencies across all ancestry informative sites with a Neanderthal allele frequency of at least 20% in at least one population. We use this subset of ancestry informative sites because we are interested in frequencies along the selected haplotype(s), which do not necessarily span the entire length of the previously described windows. If selection quickly favored introgressed alleles, we would at least expect all points in Figure 2 to show high levels of Neanderthal ancestry. However, in a number of these genomic regions, Neanderthal ancestry is almost or completely absent from East Asian and/or some ancient populations.

**Figure 2:**
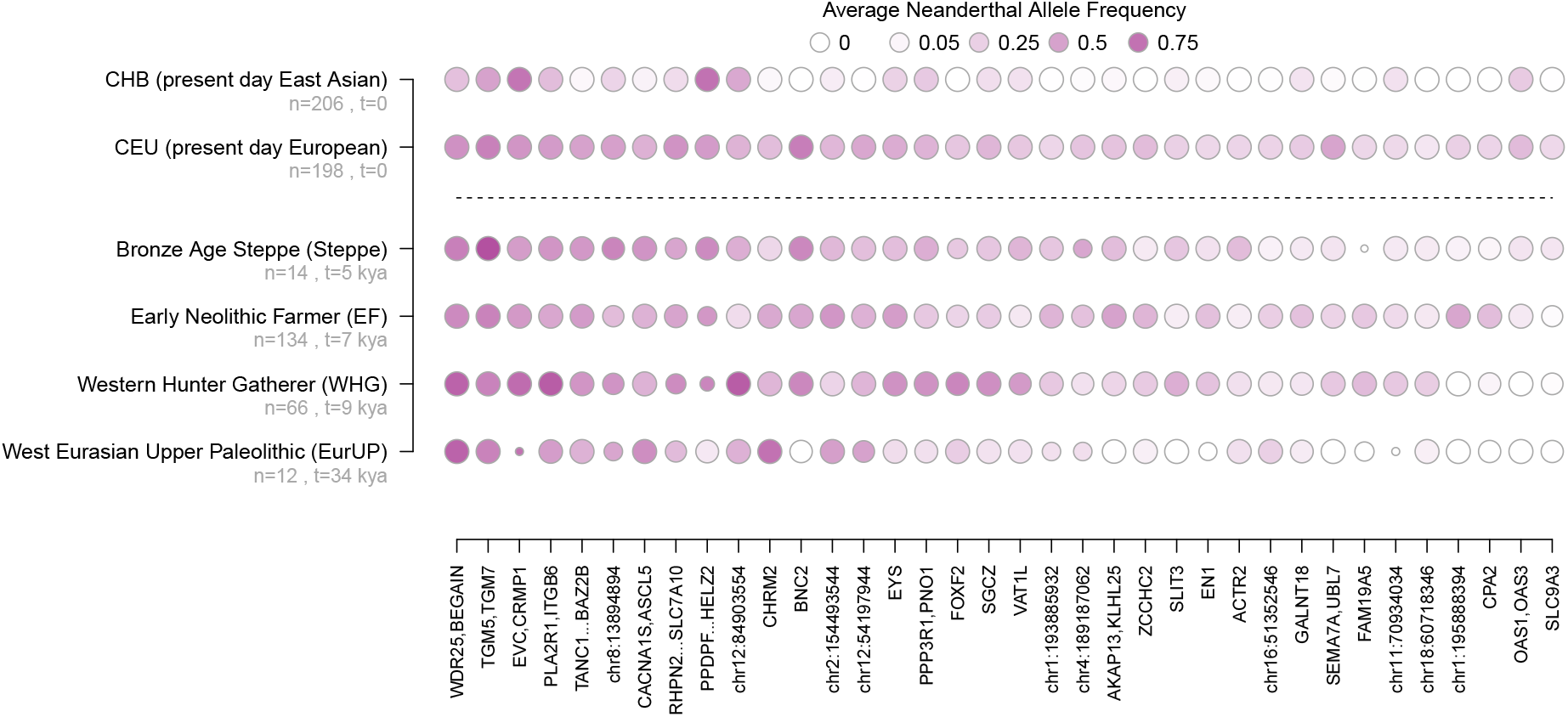
Average Neanderthal allele frequency at ancestry informative sites in ancient and present day populations in genomic regions with signatures of adaptive introgression. Darker shades of purple correspond to higher average Neanderthal allele frequencies. The size of points correspond to the number of sites over which the average Neanderthal allele frequency was calculated. Moving left to right, regions have decreasing average Neanderthal allele frequencies among ancient populations.

## 3 Method to estimate the timing of selection from introgressed haplotypes

### 3.1 Verbal Description

We first describe the intuition behind our models, before laying out the mathematical framework. We focus on how patterns of haplotype similarity among modern human populations and Neanderthals change with the start time of selection. Since we describe selection favoring a Neanderthal introgressed allele, the earliest selection can begin is the time of admixture between Neanderthals and modern humans. To accompany our description, in Figure 3 we show a cartoon of haplotype diversity and introgressed ancestry under two scenarios: immediate selection and recent selection.

**Figure 3:**
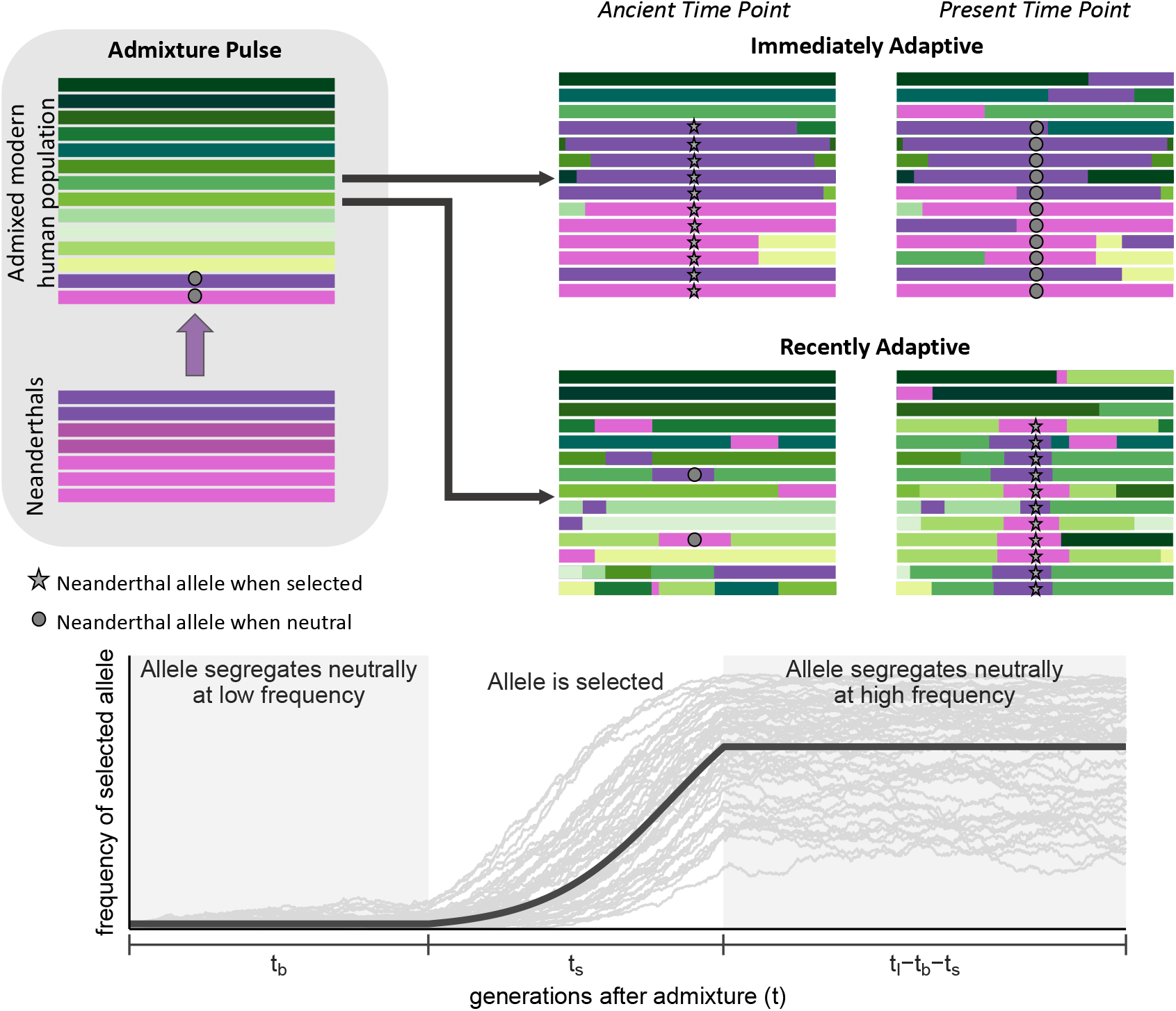
(Top) Cartoon of haplotype diversity and ancestry at three time points when selection favors Neanderthal introgressed alleles immediately (*t_b_* = 0) and when selection favors Neanderthal introgressed alleles more recently (*t_b_* > 0). Our models use a specific value of *t_b_*, but here our purpose is to visualize qualitative differences between early and recent selection. Purple tracts represent Neanderthal introgressed haplotypes, whereas green tracts represent haplotypes that did not descend from Neanderthals. We display Neanderthal ancestry tracts with fewer shades to represent their lower genetic diversity. (Bottom) Example frequency trajectory of a selected Neanderthal introgressed allele when there is a waiting time until selection. The light grey lines represent 50 stochastic trajectories in which the selected allele reached a frequency greater than 20%. The dark grey line represents the trajectory we assume in our models using the same waiting time until selection (*t_b_*), selection coefficient (*s*), and duration of the sweep phase (*t_s_*) used to generate the stochastic trajectories.

Neanderthal alleles that introgress into modern human populations start on long Neanderthal ancestry tracts (entire chromosomes at the time of admixture) that shorten over time with recombination. Since the selected Neanderthal allele takes its linked variation to high frequency with it, the timing of selection determines the genetic distance over which neutral Neanderthal alleles sweep to high frequency as well. The earlier selection is, the larger the Neanderthal haplotypes that sweep to high frequency with the selected Neanderthal allele. Conversely, the later selection is, the shorter the Neanderthal haplotypes that sweep to high frequency, such that some haplotypes in modern humans that did not introgress from Neanderthals (which we call ‘modern human haplotypes’) also rise in frequency.

Because Neanderthal genetic diversity is very low, haplotypes that introgressed from Neanderthals into modern human populations should look almost identical to each other and to haplotypes sampled from Neanderthals. Conversely, Neanderthal haplotypes are relatively distinct from modern human haplotypes compared to differences typically observed among modern human haplotypes. This is because the earliest time modern human and Neanderthal haplotypes can coalesce is when Neanderthals and modern humans shared a common ancestor, about 16,000 generations ago. Therefore, when selection brings Neanderthal haplotypes to high frequency in those modern human populations where the Neanderthal allele experienced positive selection (hereafter ‘selected populations’), selected populations gain unusually high sequence similarity with Neanderthals and unusually low sequence similarity with other modern human populations in which Neanderthal haplotypes are rare in the genomic region affected by the sweep. These patterns of unusually high and low sequence similarity persist over greater genetic distances as selection begins closer to the time of introgression.

Within selected populations, selection increases sequence similarity because the sampled haplotypes descended from one or a few ancestral haplotypes that hitchhiked to high frequency during the sweep. When selection begins earlier, these ancestral haplotypes are mainly one of the few Neanderthal haplotypes introduced by admixture. The later selection begins, the more likely these ancestral haplotypes are modern human haplotypes that became linked to the selected allele prior to the selection onset.

The putative selected Neanderthal allele did not reach fixation in the cases of adaptive introgression that we investigate, either because selection is weak and the sweep is ongoing, selection pressures changed before the Neanderthal allele reached fixation, or the phenotypic response to selection was achieved by allele frequency changes at multiple loci. Our models allow for the latter two possibilities in which selection no longer favored the Neanderthal allele, thereby potentially introducing a neutral phase after the sweep. From this phase, we can further distinguish among selection times based on how recombination distributes Neanderthal ancestry among haplotypes within the same population that do and do not carry the selected Neanderthal allele. As the partial sweep finishes, the Neanderthal alleles that hitchhike to higher frequency with the selected allele have a higher chance of recombining onto the background of the non-selected allele. The earlier the onset of selection, the more time post-sweep for these recombination events to occur, and therefore the higher the probability that alleles on the non-selected background descended from Neanderthals.

### 3.2 Model Background

Our aim is to distinguish among the possible scenarios of selection on introgressed variation. Each scenario is defined by the combination of two parameters we aim to infer in our model: the amount of time between Neanderthal admixture and the onset of selection, which we refer to as the waiting time until selection (*t_b_*) and the additive strength of selection favoring the beneficial Neanderthal allele (*s*). We build on the model-based, statistical approach introduced in Lee and Coop (2017), which uses coalescent theory to describe how different selection scenarios modify the neutral variance and covariance of population allele frequencies surrounding the selected site. This allows us to describe a multivariate normal model of population allele frequencies for each scenario, which serves as a simple approximation for their probability distribution (Weir et al., 2002; Nicholson et al., 2002).

Here, we review the framework laid out by Lee and Coop (2017). We model the change in allele frequency Δ*x_i_* = *x_i_* − *x_a_* in our sampled populations (*i*) from their common ancestral population (*a*) at the root of the tree relating all populations we consider. For neutral alleles, the population allele frequency will on average be the same as the ancestral allele frequency (E[Δ*x_i_*] = 0) because drift and hitchhiking are direction-less on average. Drift and hitchhiking do however cause an increase in the variance in the change in allele frequencies within a population, and pairs of populations can also covary in their change in allele frequency from the ancestor if they have some shared population history or gene flow, i.e. their changes in allele frequency since the ancestral population are not independent of one another. These effects are captured by the population covariance between populations, *i* and *j*, given by

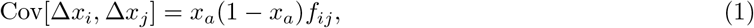

where *f_ij_* is the probability that the ancestral lineages of an allele sampled in population *i* and an allele sampled in population *j* coalesce before reaching the ancestral population (see Lee and Coop, 2017, for details).

We can estimate the neutral probabilities of coalescing among populations from neutral allele frequency data genome-wide. If we are considering *k* populations in our analysis, we thus have a *k* × *k* matrix, **F**, that describes probabilities of coalescing within and between these populations. See Appendix section A.2 for details on how we estimate this matrix. These estimated neutral probabilities of coalescing allow us to describe our neutral expectations without making any assumptions about the demographic history of populations: they implicitly account for population size, divergence, and migration. We note, however, that while this flexible approach allows for arbitrary relationships among populations, our tree requires a root that our coalescent probabilities and frequency deviations will be relative to. We define this root by placing the two most genetically distant populations on opposite sides of it. This implies that in our models, they have no shared history since the ancestral population, and therefore no chance for coalescence between the ancestral lineages of alleles sampled in each. In our analysis, the Neanderthals and Yoruba are the most distantly related, and therefore are placed on opposite sides of the root, without any subsequent gene flow. This does not negate the possibility of indirect gene flow between them; we simply define all other relationships relative to theirs.

When we incorporate the effects of selection, we model how probabilities of coalescing change with increasing genetic distance from the selected site. Importantly, all of our predictions converge to our neutral estimates because increasing recombination rates between selected and neutral sites allows for increasing independence of their coalescent history. At a far enough genetic distance, dynamics at the selected site become quickly disassociated with those at the neutral site, such that the probability of coalescing at the neutral site is that of any neutral allele unaffected by linked selection.

### 3.3 Haplotype Partitioning and the Null Model

No identified case of adaptive Neanderthal introgression has resulted in the population fixation of a Neanderthal-derived allele. These partial sweeps weaken the effect of linked selection on surrounding neutral diversity because much of the original genetic diversity prior to selection persists in these populations. In order to increase the power of our approach, we divide admixed populations according to ancestry assignments at the putative selected site. For the 1000 Genomes samples, from which we have phased haplotypes, we create two partitions: one consisting of haplotypes that carry the Neanderthal allele at the selected site (B), and the other consisting of haplotypes that carry the non-Neanderthal allele at the selected site (b). For the ancient samples, we only have unphased genotype data, and so we divide the population samples into three partitions according to individuals’ ancestry genotypes at the selected site: *BB, Bb*, and *bb*. In our models, we are concerned with a neutral allele’s ancestry background at the selected site when it is sampled. Therefore, haplotype partition *B* is equivalent to genotype partition *BB* in that any neutral allele sampled in these partitions is linked to the Neanderthal allele at the selected site. Similarly, haplotype partition *b* is equivalent to genotype partition *bb* in that any neutral allele sampled in these partitions is linked to the non-Neanderthal allele at the selected site. From here on we refer to them in our models as partition *B* or partition *b*. Predictions for genotype partition *Bb* are simply a linear combination of our predictions for the other partitions, which we describe in Supplement section S3.5.

Our haplotype partitioning of the data requires us to modify our null model from that of Lee and Coop (2017), who focused on full sweeps and simply used the genome-wide neutral **F** matrix to parameterize their null model. The neutral probabilities of coalescing that we estimate between any pair of populations (*i, j*) are essentially averaged over all possible migration histories of our alleles. However, our population partitioning scheme alters the probability of each migration history because we split populations based on ancestry at the partition site. When we create a population that only carries Neanderthal alleles at the partition site (partition *B*), nearby sites will have a much higher frequency of Neanderthal alleles relative to the full population. Therefore, even under neutrality, the probability that a randomly sampled allele in partition *B* will coalesce with an allele sampled in Neanderthals is much higher than in the full population case.

Under population partitioning, we can describe our null coalescent probabilities as a function of the genome-wide neutral probabilities of coalescing and the recombination rate (*r*) between the partition site and neutral site of interest. We first consider the relationship between Neanderthal population *n* and partition *B* of population *i* in which all haplotypes carry a Neanderthal allele at the partition site. We define *t_I_* as the number of generations ago that Neanderthal alleles introgressed into modern human populations. We are interested in the probability that the neutral lineage remains linked to the same Neanderthal allele at the partition site that it was linked to at sampling by the time of admixture. We can approximate this probability as *e^−rt_I_^*. By remaining linked to the Neanderthal allele at the partition site, we know that it, too, introgressed from Neanderthals and thus will coalesce with the Neanderthal allele at the same site with approximately the same probability as any two alleles sampled from Neanderthals (*f_nn_*). If it does recombine out at some point before admixture, it has the population’s neutral probability of coalescing with Neanderthals. This neutral probability of coalescing still accounts for the possibility that the neutral allele descended from Neanderthals, when the average Neanderthal-introgressed allele frequency is the initial admixture fraction (*g*). We define probabilities of coalescing under the null model as 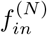. In all we derive the probability of coalescing between a pair of lineages sampled from partition *B* of population *i* and Neanderthal population *n* to be

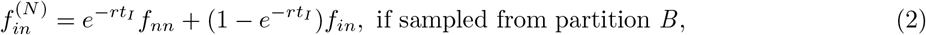

where superscript (*N*) refers to predictions under our null model. In partition *b* of the same population (*i*), all alleles are linked to a non-Neanderthal allele at sampling. If an allele sampled in this population never recombines out, we know it cannot have introgressed. Therefore in order to have the chance at coalescing with a Neanderthal allele, it must recombine out before admixture. Forward in time, that means that a Neanderthal allele recombined onto the sequence carrying the non-Neanderthal allele at the partition site. Therefore a neutral allele sampled from partition *b* of population *i* has the following probability of coalescing with a neutral allele sampled from Neanderthals:

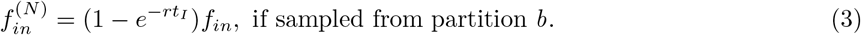

### 3.4 Selection Model

Selection favoring Neanderthal alleles in modern humans modifies probabilities of coalescing from neutral expectations because it increases levels of Neanderthal ancestry around the selected site in selected populations. Thus, in the whole (non-partitioned) population, the probability that an allele descended from Neanderthals is much higher than the original admixture proportion, as we described informally above in our discussion of Figure 3.

#### 3.4.1 Derivation

In selected populations, we consider three phases in the selected allele frequency trajectory: the neutral phase following admixture in which the Neanderthal-derived variant is at frequency *g* for *t_b_* generations (neutral phase I), the sweep phase in which the variant rises from frequency *g* to frequency *x_s_* in time *t_s_*, and the neutral phase between the sweep finish and the present in which the variant remains at frequency *x_s_* (neutral phase II). When the relative fitness advantage of the selected allele is additive, such that heterozygotes have an advantage of s and homozygotes 2s,

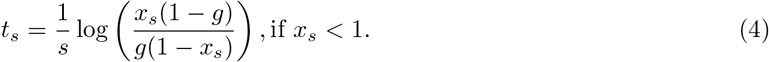

Since the sum of all three phase durations equals the time between the present day and admixture (*t_I_*), the duration of neutral phase II equals *t_I_ − t_b_* − *t_s_*.

As the waiting time until selection *t_b_* increases, and assuming the same *t_b_* for all selected populations, we transition from describing a case in which the selected allele became beneficial in the common ancestor of all Eurasian populations, to the case in which the selected allele became beneficial independently in each selected population. If *t_b_* is short enough such that selection began in the common ancestor of a group of selected populations, we must account for the possibility that this common ancestor also has descendent populations that do not carry the selected allele at high frequency, possibly due to subsequent drift or negative selection. According to *t_b_*, we assign each population with very low frequency of the selected allele into one of two categories: (i) ancestors never selected or (ii) ancestors selected with subsequent loss of the selected allele in this population. We assign the first category if the low frequency population does not share a common ancestor with any selected populations at the time selection starts, i.e. its divergence from all selected populations predates selection. Otherwise we assign the second category.

Similar to the null model, we condition on ancestry at the putative selected site and consider how recombination modulates ancestry with increasing genetic distance. The most important difference between the null and selection model is neutral phase II: in the selection model, if a neutral lineage recombines, it has a high probability of recombining onto the selected allele’s background. Therefore, there is a higher chance that the neutral allele itself descended from Neanderthals.

##### Between Neanderthals and selected populations

In this section we derive the probabilities of coalescing between the haplotypes in each partition of selected human populations and Neanderthals. When we sample an allele, we know whether or not it is linked to the selected Neanderthal allele based on the partition it belongs to. Conditioning on this ancestry background at sampling, we focus on whether the ancestral lineage is linked to the selected Neanderthal allele when we transition between each phase looking backwards from the present to admixture.

First, we determine the probability that a neutral lineage is linked to the selected Neanderthal allele at the transition point between neutral phase II and the sweep completion (which occurs at time *t_I_* − *t_b_* − *t_s_*). If there is never a recombination event between the selected and neutral site during neutral phase II, then the neutral lineage remains associated with its initial ancestry background at the time of the sweep completion. Alternatively, if at least one recombination event occurs, then the final recombination event determines the ancestry background that the neutral lineage is associated with at the time of the sweep completion. The frequency of the selected allele determines the probability that a recombining neutral lineage becomes associated with the selected background. Thus with probability *x_s_*, the neutral lineage becomes associated with the selected Neanderthal allele, and with probability 1 − *x_s_* it becomes associated with the non-Neanderthal allele at the selected site. Therefore, the probability that a neutral lineage sampled from a selected population is linked to the selected Neanderthal allele at the transition point between neutral phase II and the sweep completion is

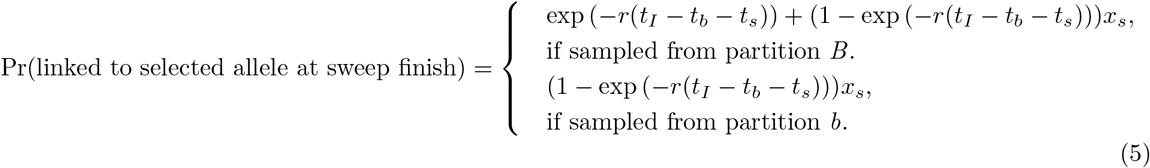

in which an allele sampled from partition *b* can only be linked if it recombines off of its background.

Second, during the sweep phase, an allele that begins linked to the selected allele will always be associated with that background with probability

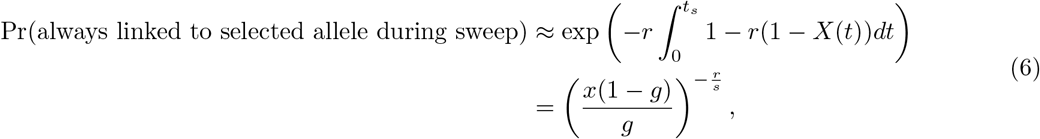

where *X*(*t*) is the frequency of the selected allele in generation *t* of the sweep, following a deterministic, logistic trajectory. In words, this probability is approximately the product of the probabilities of not recombining onto the non-selected background each generation of the sweep. Similarly, an allele that begins the sweep phase linked to the non-selected allele will always be associated with that background during the sweep with probability

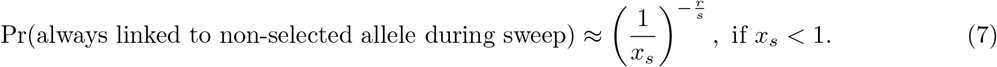

The above associations persist if the lineage never recombines out of its background during neutral phase I, with approximate probability *e^−rt_b_^*. So, if a neutral lineage remains linked to the selected allele during the sweep phase and fails to recombine during neutral phase I, it must have descended from Neanderthals and thus coalesces with the lineage sampled from Neanderthals with approximately the same probability as any two lineages sampled from Neanderthals, *f_nn_*. If the neutral lineage remains linked to the non-selected allele throughout the sweep phase and fails to recombine during neutral phase I, it definitely did not descend from Neanderthals and thus cannot coalesce with an allele sampled from Neanderthals. If a lineage becomes disassociated with its background at some point during the sweep phase, and/or recombines out at least once during neutral phase I, we assume it has its population’s neutral probability of coalescing with Neanderthals. Our approximation ignores the possibility that a lineage linked to the non-selected allele can recombine onto the selected allele’s background during the sweep. In total, the ancestral lineage of an allele sampled from an admixed, selected population coalesces with the ancestral lineage of an allele sampled from Neanderthals with probability

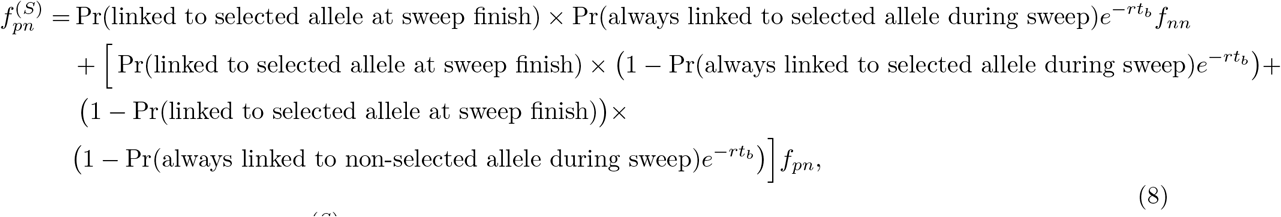

where the superscript *S* in 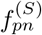 refers to our predictions under the selection model, and subscript *p* denotes any partition of any selected population. In Supplement sections S3.1 and S3.2 we illustrate predictions for other population relationships under the selection model. We follow by describing modifications under both the null and selection models to incorporate ancient samples (section S3.3), migration among admixed modern human populations (section S3.4), and genotype partition Bb (section S3.5). Our approximations provide a good fit to those obtained by simulations, see Figure 4.

**Figure 4:**
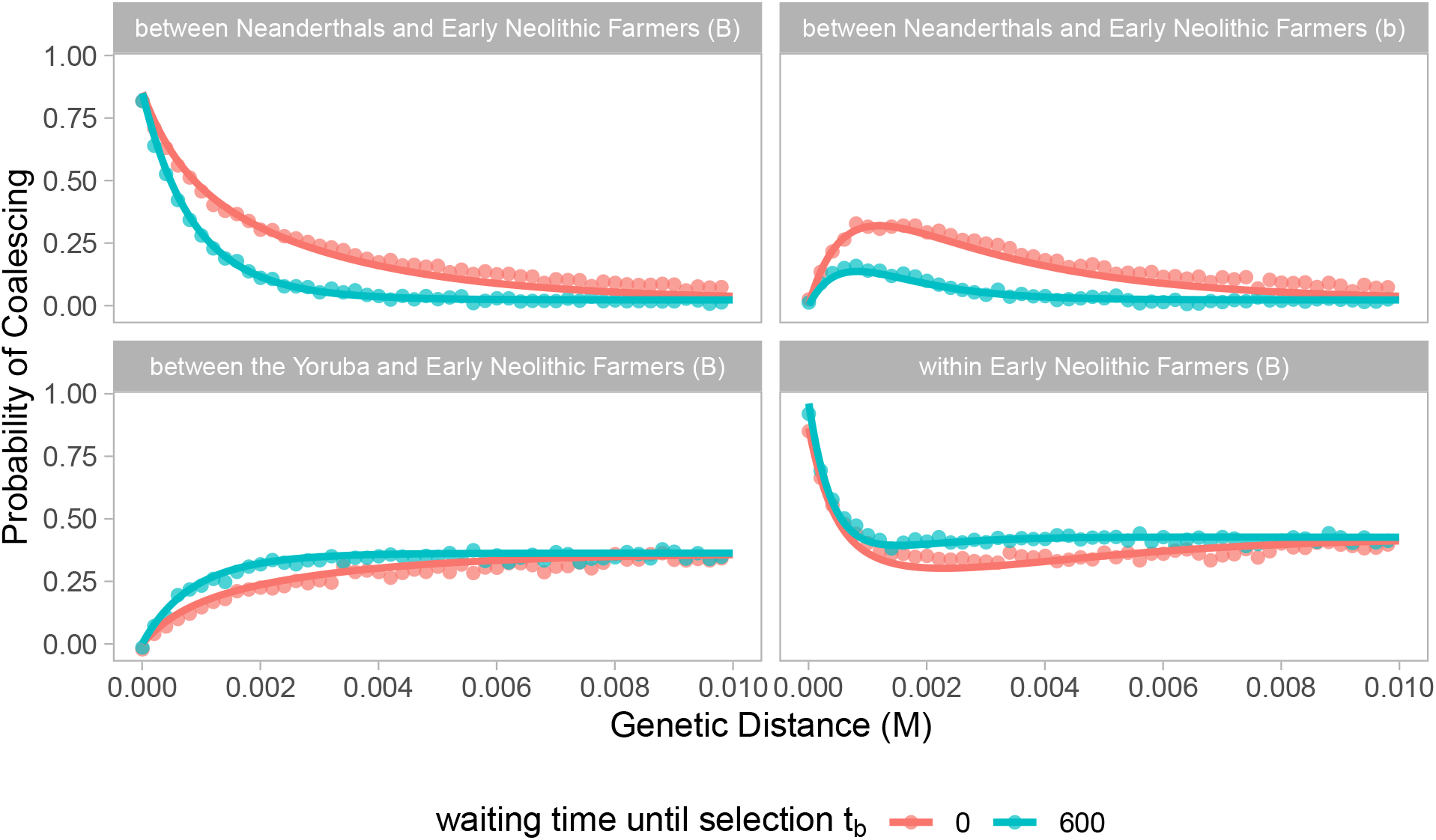
Probabilities of coalescing between pairs of alleles at increasing genetic distances from the selected site. Analytical predictions (lines) closely match estimates averaged across 250 simulations (points). Here, we show results for a case in which the Neanderthal derived allele became adaptive in the ancestors of Early Neolithic Farmers immediately upon introduction (*t_b_*=0, red) and after 600 generations (blue). The favorable allele experienced a selective advantage (*s*) of 0.01 and on average reached a frequency (*x_s_*) of 0.7 before becoming neutral again. The genetic window corresponds to 1 Mb with a constant recombination rate of 10^−8^. Partition *B* refers to alleles linked to the beneficial Neanderthal allele, while partition *b* refers to alleles linked to the non-Neanderthal counterpart at the selected site. We present categories of population relationships in which probabilities of coalescing vary the most with different waiting times until selection. See Figure S12 and Figure S13 for more population relationships, waiting times until selection, and final frequencies of the selected allele.

### 3.5 Inference

For each model (null or selection) and combination of free parameters, we define a variance-covariance matrix of population allele frequencies for a given distance away from the selected site (Lee and Coop, 2017). We approximate the joint probability distribution of population allele frequencies at a neutral locus (*l*) as being multivariate normal around the ancestral allele frequency (*x_al_*) with covariance equal to *x_al_*(1 − *x_al_*) times the modified matrix of coalescent probabilities (**F**^(**N**)^ or **F**^(**S**)^) (Weir et al., 2002; Nicholson et al., 2002; Samanta et al., 2009; Coop et al., 2010). In both the null and selection models, the modified matrices of coalescent probabilities depend on the genetic distance to the selected site (*r_l_*), the genome-wide neutral matrix of coalescent probabilities (**F**), and all of the admixture graph parameters of interest (*A_G_*), which describe admixture proportions and timing among Neanderthal and admixed modern human populations as well as divergence and sampling times of admixed modern human populations (see Figure 1 and Table S2). The selection model’s **F**^(**S**)^ depends on three additional parameters: the strength of selection (*s*), the waiting time until selection (*t_b_*), and the final frequency of the selected allele (*x_s_*). Therefore at a neutral polymorphic site with allele frequency data in our populations, we can estimate the probability of the observed population allele frequencies 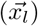 under the selection model as

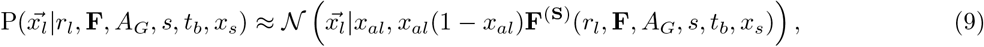

where the set of parentheses following **F**^(**S**)^ denote the parameters of the function **F**^(**S**)^. When we predict **F**^(**S**)^, we categorize Neanderthal-admixed populations as ‘selected’ if their frequency of the Neanderthal allele at the selected site is greater than or equal to 0.05. We set *x_s_* to be the average selected Neanderthal allele frequency among all of these putative selected populations. Our method could be extended to allow *x_s_* to vary among populations and to do model choice of the selected populations, however for simplicity we do not pursue those applications here. We estimate the composite likelihood of free parameters s and *t_b_* given the allele frequency data across populations (*D*) in a window around a candidate selected site by taking the product of all likelihoods calculated at each locus to the left and right of the candidate selected site as follows,

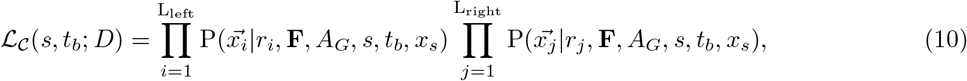

where L_left_ and L_right_ represent the number of polymorphic loci to the left and right of the candidate selected site. We calculate composite likelihoods among the proposed parameter combinations of *s* and *t_b_* shown in Table S4.

The composite likelihood under the null model takes the same form as selection; we simply replace **F**^(**S**)^ with **F**^(**N**)^ and remove dependence on s, *t_b_*, and *x_s_*. Each partition site represents a potential selected site and thus differs in its genetic distance to each neutral locus. Therefore, each partition site has its own set of composite likelihoods for the null and selection models. We identify the ‘best’ partition site for each region by selecting the site whose ratio of its maximum composite likelihood under the selection model to the null model is greatest, i.e. by maximizing the level of support for selection at this site. From this partition site, we identify the maximum composite likelihood estimates 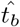 and 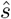 for the region. We note that in our application, these estimates tended to be very similar among partition sites.

Our method focuses on inferring the history of selection from haplotypes alone, and does not incorporate the distribution of the selected allele frequency across populations (as in Racimo et al., 2018; Refoyo-Martínez et al., 2019). We could incorporate into our composite likelihoods the conditional probability of *x_s_* across populations under a given parameterization of the selection model, but here our focus is on the information about the timing of selection contained in haplotype patterns.

We analyze bi-allelic sites that are polymorphic among our set of samples. Since our multivariate normal approximation works best when alleles segregate ancestrally at some intermediate frequency, we remove sites polymorphic in only one population with a minor allele frequency less than 0.01. In practice, we mean centered our observed allele frequencies, which removes dependence on the ancestral allele frequency *x_al_*. We provide more detail on mean centering, sample size correction, and implementation in Appendix section A.3.

#### Distinguishing between immediate and non-immediate selection

To assess the timing of selection relative to introgression we first distinguish between immediate (*t_b_* = 0) and non-immediate (*t_b_* > 0) selection. We do so by using the ratio of the maximum composite likelihood under all parameter combinations to the maximum composite likelihood when selection is immediate,

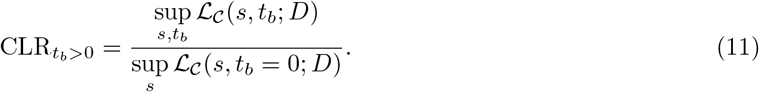

The higher this ratio, the more support we have in favor of an initial neutral period relative to immediate selection. Since composite likelihoods ignore the correlation in allele frequencies across loci due to linkage disequilibrium, we cannot use traditional statistical methods for model selection. Therefore we rely on simulations and reject the immediate selection case if the upper 95th percentile of CLR_*t*_*b*_>0_ from immediate selection simulations does not exceed the observed value in a region. In other words, we reject the null hypothesis of immediate selection if we observe a value of CLR_*t*_*b*_>0_ high enough to be unlikely if selection were truly immediate. The following sections contain more detail on the simulation procedure used to reject immediate selection.

#### Validation

We ran our method on simulated data to evaluate its performance, using SLiM 3.0 for forward in time simulations (Haller and Messer, 2019). We simulated 2cM (2 Mb) loci with the selected mutation at the center of the locus. We simulated under the demographic history shown in Figure 1, along with a divergence time between the Yoruba and Eurasian populations of 2,500 generations and a divergence time between modern humans and Neanderthals of 16,000 generations. The Neanderthal population was simulated with a population size of 3,000, whereas all others were simulated with a population size of 10,000.

Our SLiM simulations generated tree sequences, to which we added neutral mutations and calculated population allele frequencies using the same sample sizes as in our real data in Python version 3.7.4 with `msprime` (Kelleher et al., 2016), `tskit` (Kelleher et al., 2018), and `pyslim` (Haller et al., 2019). We ran our method on these data in R version 3.4.4. The method assumed the same demographic history that we simulated and used a neutral **F** estimated from neutral allele frequency data, also produced by SLiM simulations under the same demographic history as the selection simulations. See Appendix A.1 for more simulation details.

Our first goal was to determine our power to detect selection that did not begin immediately, and so we established the significance cutoff using simulations under the null model of immediate selection. Running the method on simulations of immediate selection, we identified a positive correlation between xs, the average selected allele frequency among all putative selected populations, and CLR_*t*_*b*_>0_, the composite likelihood ratio quantifying the method’s support for non-immediate selection relative to immediate selection (Figure S1). This relationship between xs and CLR_*t*_*b*_>0_ among our immediate selection simulations does not differ among combinations of the selection coefficient (*s*) and the duration of the sweep (*t_s_*) (Figure S1). Thus, for each simulation of non-immediate selection, we rejected immediate selection if its CLR_*t*_*b*_>0_ was greater than the upper 97.5th percentile of CLR_*t*_*b*_>0_ from the 500 immediate selection simulations with the closest *x_s_*. We used a similar procedure to determine the significance cutoffs for the real dataset.

Varying the onset time of selection in our simulations, we found that the method’s power to reject immediate selection increases with the true waiting time until selection (*t_b_*) and *x_s_* (Figure 5). Our method has reasonable power to reject immediate selection when selection begins > 1000 generations after admixture, suggesting that we should correctly identify many of the regions that only recently contributed to adaptation. Overall, s is not well inferred, likely because we deal with old selection events. Therefore, while s appears in our selection model, we concentrate on inferring *t_b_* in our application. The method yields somewhat biased estimates of the waiting time until selection (Figures S2–S5). However, among the simulations in which the method rejected immediate selection, the method does a good job tracking the true time until selection (Figure 6). This performance decreases when fewer populations are considered to be selected, but still the method can identify that selection started much later in time. For more details on method performance, see Supplement section S2.1.

**Figure 5:**
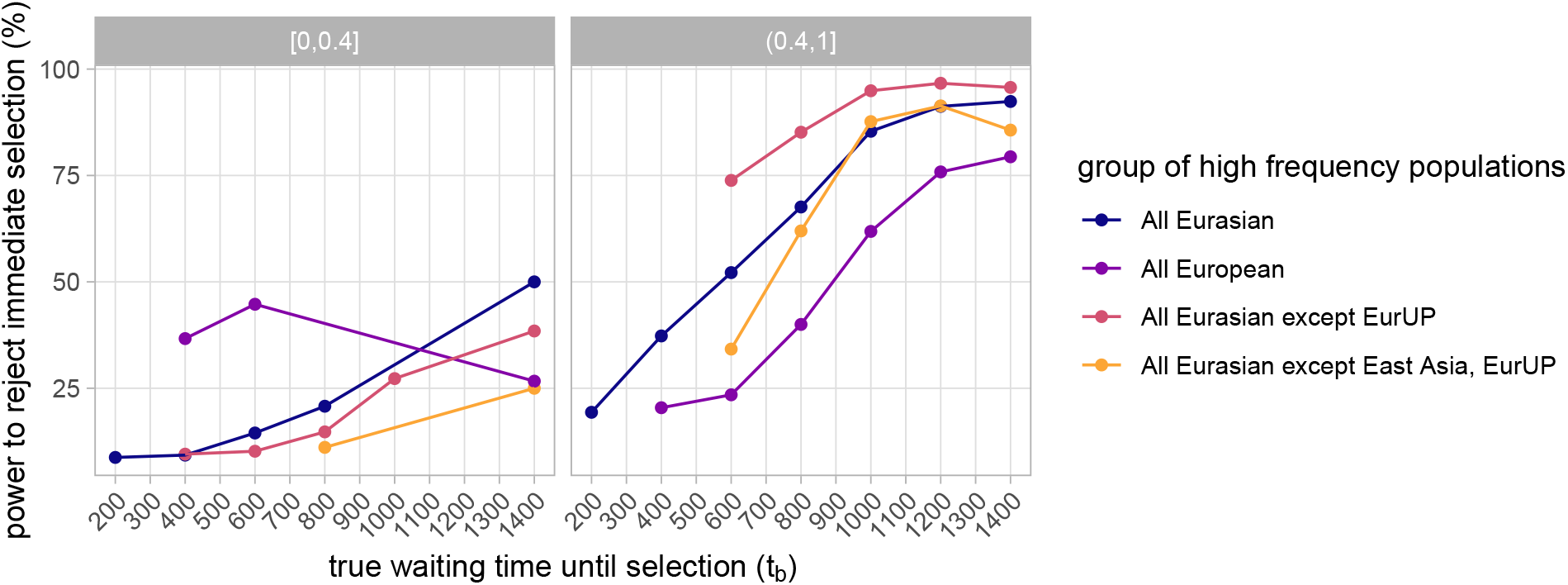
Power to reject immediate selection as the true waiting time until selection (*t_b_*) increases. Lines correspond to simulations with different groups of high frequency populations (each with the selected allele frequency greater than 0.05), i.e. groups of populations considered selected. Note that this does not perfectly correspond to groups of populations truly selected in our simulations, and so we have different numbers of observations contributing to each point. Points with fewer than 10 observations were removed. Panels correspond to bins of *x_s_*, the selected allele’s average frequency among populations considered selected. Results for *x_s_* > 0.4 are qualitatively similar, though we show finer bins in Figure S7.

**Figure 6:**
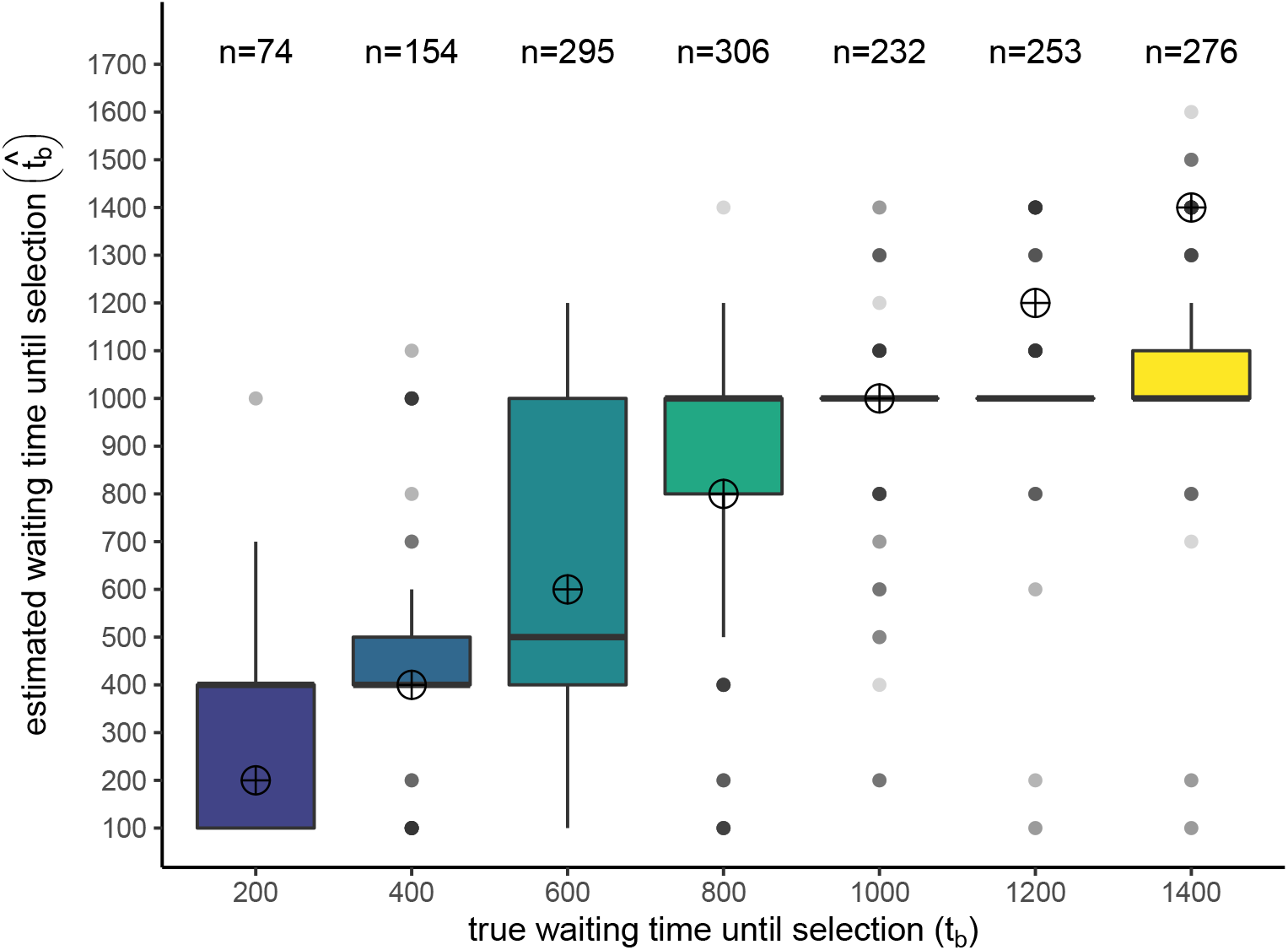
Estimated waiting time until selection 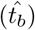 among simulations with all populations considered selected (each *x_s_* > 0.05) and in which the method rejected immediate selection. Target points mark the true waiting time until selection that we aim to estimate. Earlier true waiting times have fewer observations because the method has less power to reject immediate selection in this parameter space. Results for more groups of putative selected populations are shown in Figure S6. Estimated waiting times until selection under the immediate selection case can be found in Figure S1.

## 4 Data Availability

Scripts for imputation, simulations, and the method are provided at http://www.github.com/SivanYair/selTime_neanderthal_AI. The script that computes genotype likelihoods can be found at https://github.com/mathii/gdc3/blob/master/apulldown.py. File Supp_Table_1.csv contains Table S1, available at FigShare.

## 5 Data application to estimate the timing of selection on Neanderthal alleles

### Procedure for identifying selected site and parameter estimates

For each region, we ran our method on the previously defined windows in addition to 1 cM flanking them on each side, because if selection is immediate with *s* = 0. 01 it takes about 1 cM for the signals of selection to almost completely decay. We used the HapMap Project’s genetic map to identify the endpoints of genomic analysis windows and recombination rates between each neutral and selected site for the method (note that this means that we need to assume that recombination rates have been relatively constant over this time period). We excluded sites without Neanderthal allele frequency data from our analysis, as they are less informative of introgression patterns and the method might instead pick up on unrelated signals of selection at these sites.

We estimated the neutral **F** from putative neutral allele frequency data genome wide. We chose regions at least 300 base pairs long, at least 0.4 cM away from a gene, and with a minimum per base pair recombination rate of 0.9 × 10^−8^ using the ‘Neutral Regions Explorer’ (Arbiza et al., 2012). This led to 105,786 sites after filtering. While these sites may not be entirely unaffected by linked selection, we can still use them to represent background patterns of genetic diversity from which we distinguish strongly altered patterns caused by adaptive introgression.

Among the ancestry informative sites in each region, we chose partition sites (which represent potential selected sites) to be all sites with at least two ancient populations with data and a Neanderthal allele frequency ≥ 20% in at least one European, East Asian, or South Asian 1000 Genomes population. For a region, we looped over possible partition sites to run the method on their corresponding dataset, and then selected among these partition sites. Then from the ‘best’ partition site we selected among parameter estimates of *t_b_* and *s*. For example, in Figure 7 we show the profile composite likelihood surface of *t_b_* at the maximum composite likelihood partition site (relative to the null model at this site) in the region OAS1,OAS3. The peak in this surface corresponds to our estimate of *t_b_*, which in this region we found to be very high. As the relationships among our populations are only partially understood, we ran the method with the Altai Neanderthal sample included in the Neanderthal population (discussed in Supplement section S2.2.1 and Figure S10) and with multiple plausible modifications to the admixture graph in Figure 1 (discussed in Supplement section S2.2.2 and Figure S11). To account for the uncertainty in our imputation procedure, in each region we reran the method 40 times on bootstrapped datasets: rather than using the maximum likelihood genotype at the partition site to assign an ancient sample to a genotype partition, in each bootstrap run we randomly assigned the sample to one of the partitions according to its posterior genotype probabilities provided by the imputation algorithm (results shown in Figure S9).

**Figure 7:**
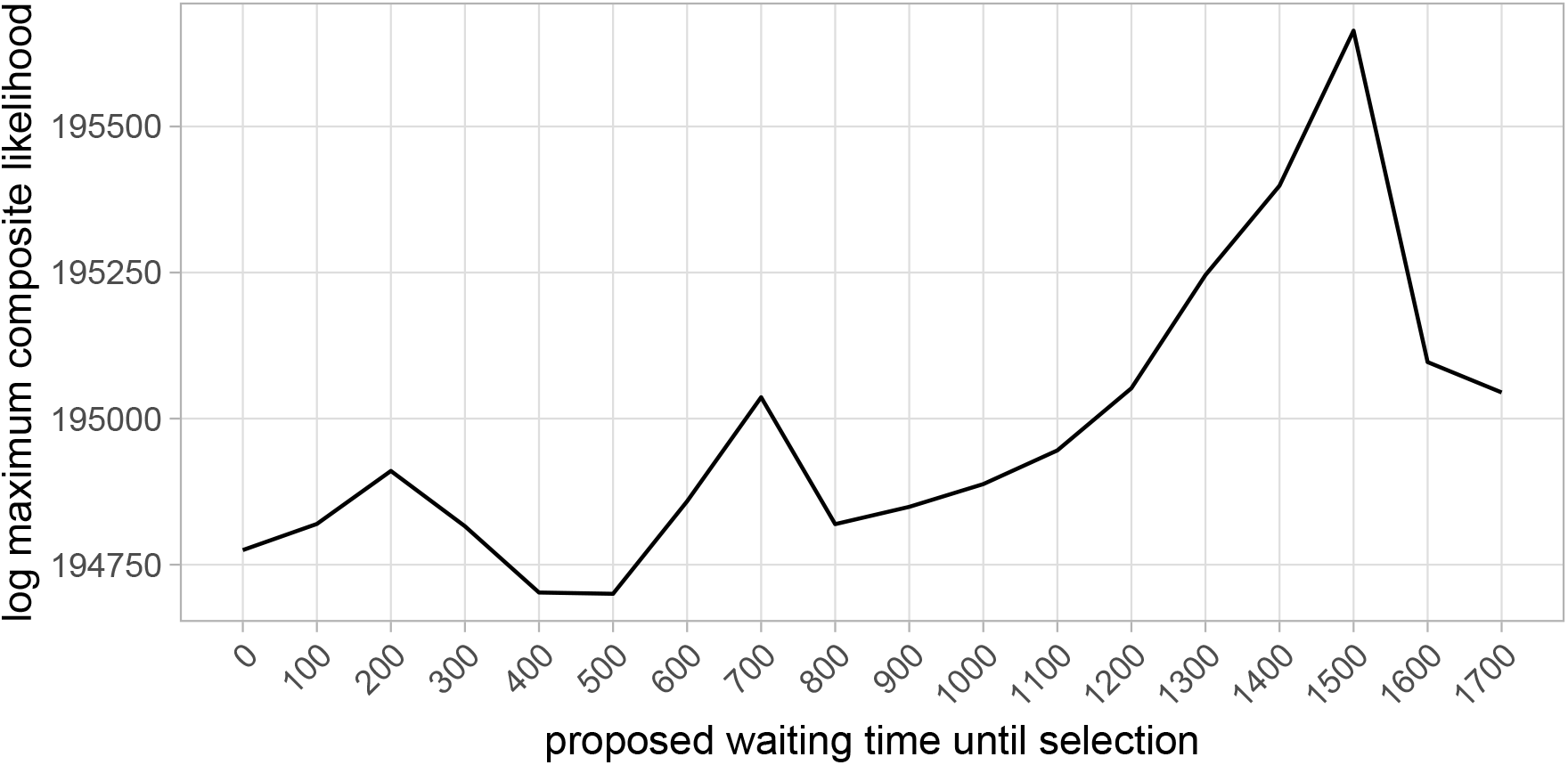
Profile composite likelihood surface of the waiting time until selection (*t_b_*) in the region OAS1,OAS3.

### Procedure for distinguishing between immediate and non-immediate selection

Among the regions in which the method estimated a non-immediate waiting time until selection 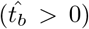, we rejected immediate selection if their CLR_*t*_*b*_>0_ exceeded the upper 97.5th percentile of those from the 500 immediate selection simulations with the closest xs (the average selected allele frequency among all putative selected populations). Recall that we can only identify these significance thresholds with simulations. Since composite likelihoods may be sensitive to the spatial distribution of analyzed sites and their corresponding populations with known allele frequencies, we sampled sites to analyze in the simulated data using the same spatial patterning as the region of interest. Specifically, we binned the genetic positions of analyzed sites into 10^−3^ cM intervals to the left and right of the selected site. For each of these sites, we sampled a site in the simulated data from the same genetic distance bin and masked its allele frequencies from populations with no data. We used the same immediate selection simulations as in the Validation section, except this time evaluating them over a larger 3 Mb (3 cM) locus.

Of the 36 regions we ran our method on, our method estimated 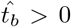 in 26, and rejected immediate selection in 17 (Figure 8). Four of the regions in which we rejected immediate selection had a standard deviation greater than 100 generations of 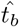 in their bootstraps over the imputation uncertainty: CHRM2, chr1:193885932, chr2:154493544, and chr12:84903554 (Figure S9). For the remaining 13 regions in which we have greater confidence in our estimated waiting times, nine of these regions have 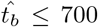, whereas the remaining four have extremely high estimates: SEMA7A,UBL7 has 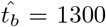 and CACNA1S,ASCL5; OAS1,OAS3; and chr4:189187062 have 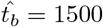.

**Figure 8:**
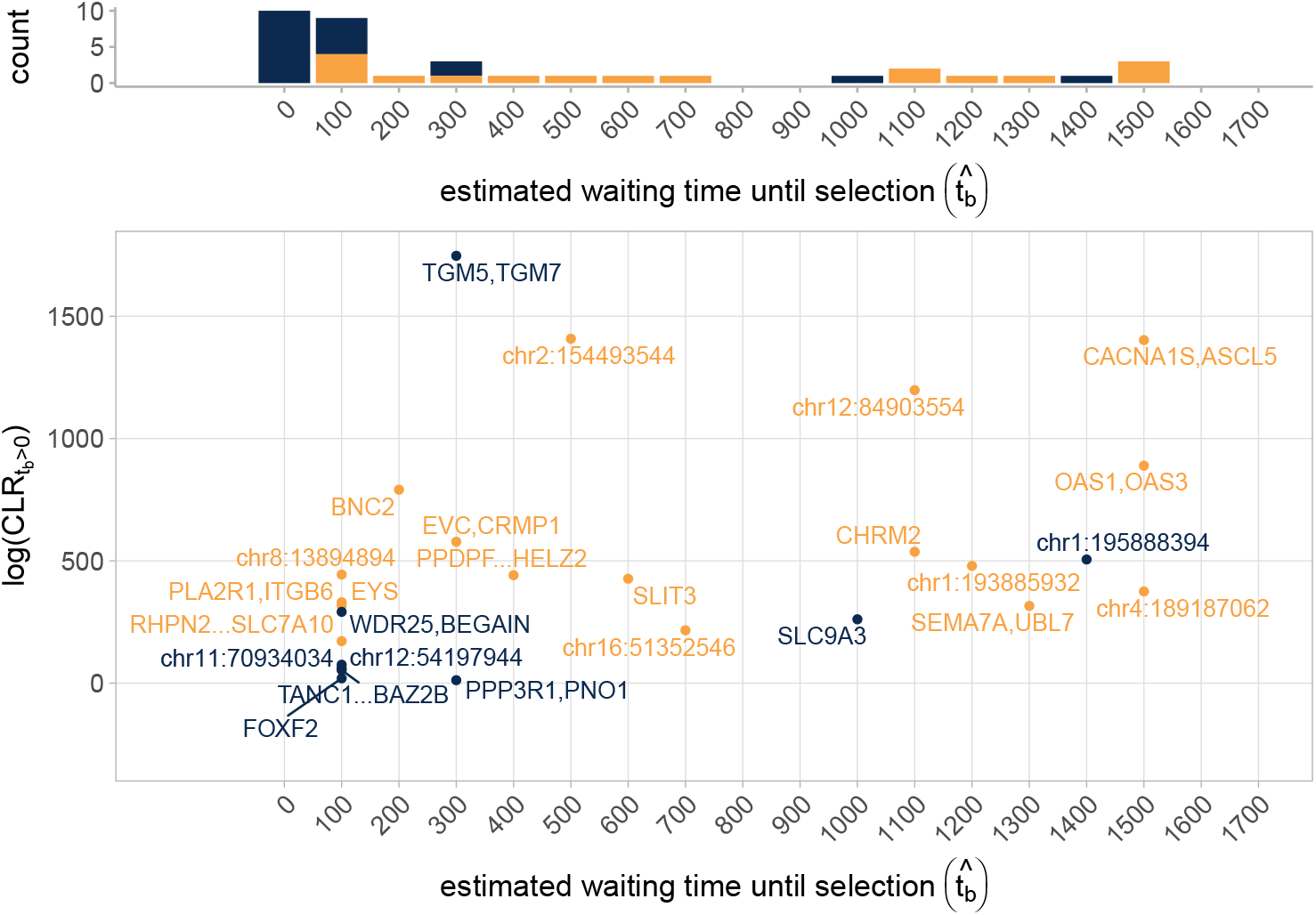
(Top) Distribution of estimated waiting times until selection 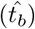 in the application of the method to 36 regions of adaptive introgression. Blue bars represent cases in which the method did not reject immediate selection, and orange bars represent cases in which the method did reject immediate selection. Blue bars are stacked on top of orange bars (they do not overlap). (Bottom) Results among all regions with 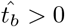. These represent 26 out of 36 cases. Regions in which the method did not reject immediate selection are colored blue, and regions in which the method rejected immediate selection are colored orange (17 cases).

As with many selection scans, the function of alleles underlying adaptation are obviously unknown. Among the set of regions with a shorter initial neutral period, BNC2 is a better-studied candidate of adaptive Neanderthal introgression, where the likely selected Neanderthal variant is associated with lighter skin pigmentation (Dannemann et al., 2017). As for the other regions with early selection, mutations in EYS affect the development and maintenance of photoreceptors in the retina (Collin et al., 2008; Abd El-Aziz et al., 2008; Alfano et al., 2016), proteins encoded by EVC are involved in the regulation of Hedgehog signaling and mutations in this gene affect patterning during development (Ruiz-Perez et al., 2000; Blair et al., 2011; Caparrós-Martín et al., 2013; Pusapati et al., 2014), proteins encoded by CRMP1 may be involved in the development of the nervous system and epithelial sheets (Nakamura et al., 2014; Yu-Kemp et al., 2017), and SLC7A10 contributes to synaptic regulation in the central nervous system (Ehmsen et al., 2016; Palaćin et al., 2016; Mesuret et al., 2018). HELZ2 is involved in regulating the differentiation of adipocytes and could thus play a role in metabolism (Katano-Toki et al., 2013). PLA2R1, ITGB6, and SLIT3 putatively play a role in tumor suppression, however mutations at all of these loci are also known to influence variation in other phenotypes (Marlow et al., 2008; Hezel et al., 2012; Guo et al., 2013; Bernard and Vindrieux, 2014; Yu et al., 2014; Ng et al., 2018; Yoshikawa and Asaba, 2020). Most research on RHPN2 and PTK6 has focused on the role of their expression in cancer progression (e.g. Zheng et al., 2010; He et al., 2015). The remainder of regions in this category are poorly studied or intergenic and far from genes. As for the recently selected regions, SEMA7A plays a role in both the immune and nervous systems, regulating active T cells in the former and axon growth and formation during development in the latter (Pasterkamp et al., 2003; Suzuki et al., 2007; Gras et al., 2013). Polymorphisms at this locus are also associated with variation in bone mineral density (Koh et al., 2006). UBL7 may play a role in ubiquitin signaling, however little is known about the effects of variation at this locus (Zhang et al., 2017). CACNA1S encodes a protein known to contribute to the structure of calcium channels involved in skeletal muscle contraction (Tanabe et al., 1990; Ptáček et al., 1994; Ertel et al., 2000) and in some cases can influence body heat regulation (Beam et al., 2017). ASCL5 is less well studied, though mutations in this gene have been implicated to affect tooth development (He et al., 2019). OAS1 and OAS3 haplotypes can influence innate immunity, which we discuss in more detail later. Based on what we currently understand about mutations in these genomic regions, the non-immediate benefits provided by Neanderthal alleles may have varied widely and were perhaps associated with different environmental factors.

## 6 Discussion

Here we use patterns of linked selection to develop one of the first model-based methods that infers the time at which Neanderthal introgressed alleles became adaptive. Our approach uses ancient DNA as well as the hitchhiking effect to investigate the temporal history of selection. We directly address partial sweeps, which likely reflect most cases of human adaptation and perhaps adaptive introgression more generally, and use them as a tool to time the onset of selection. While we require a known demographic history among Neanderthal-admixed populations, our method is robust to modifications in these assumptions (see Supplement section S2.2 for discussion).

We found that in most regions that we analyzed, we could not reject a model of selection immediately favoring Neanderthal-introgressed alleles upon admixture. In some of these regions, East Asians and/or ancient populations do not contain signatures of adaptive introgression. Obviously some of these cases may represent our lack of power to rule out short onset times from haplotypes alone and so our results should not be taken as indicating that selection began immediately for many Neanderthal haplotypes. In addition, the regions that we ran our method on were identified from signals of unusual sharing with Neanderthals (Racimo et al., 2017), and so we may be missing some cases of recent selection on Neanderthal introgression that dragged up only very short blocks of Neanderthal ancestry. We also limited our analysis to regions with signatures of adaptive introgression in European populations, so there may be cases of recent selection in East Asian or other populations that we did not identify. Among the regions in which we did reject immediate selection, we estimated a wide range of waiting times until selection. From simulations, we found that we can generally classify these cases into earlier 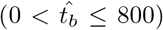 and more recent 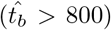 selection. These time frames may be coarse, but with our estimates of recent selection being far from the boundary between early and recent selection, the transition between them possibly marks the end of the last glacial period (~ 11.5 kya). Thus, we are potentially distinguishing between adaptation to conditions during the Upper Paleolithic versus later periods defined by warming, technological innovation, the emergence of farming, and higher population densities.

We modeled that the beneficial allele was at a low frequency for some time followed by the onset of selection due to some environmental change. This initial period could correspond to the Neanderthal allele being neutral or balanced at low frequency. However there are a few possibilities other than a truly delayed onset of selection that could explain our rejection of immediate selection. First, selection may have immediately favored an allele, but the allele was recessive and thus took a long time to rise in frequency. We chose to model additive selection because we do not have information on dominance, but for recessive alleles we may be timing its rise to intermediate frequency rather than the selection onset (see Jones et al., 2020, for a possible dominance related lag in an introgressed pigmentation allele in snowshoe hares). Given that the method does a poor job estimating the selection coefficient for these old sweeps from haplotype data alone, we likely cannot distinguish among different models of dominance. Second, selection may have immediately favored a beneficial Neanderthal allele, but it was only able to rise in frequency once it recombined off other tightly linked deleterious alleles on its Neanderthal haplotype background. There are two ways that this could lead us to incorrectly infer more recent selection: (*i*) the beneficial allele could have recombined earlier than expected, causing it to rise in frequency close to its introgression time but on a shorter Neanderthal haplotype than we predict under immediate selection, or (*ii*) a time lag between the selection onset and the recombination event that allowed the allele to rise in frequency could cause our timing estimates to mark the recombination event rather than the selection onset. Some studies have described the conditions in which an adaptive allele could introgress and eventually fix given its linked deleterious background upon introduction (e.g. Uecker et al., 2015), with some investigation into the timescale in which recombination would generate a haplotype with a net selective advantage (Sachdeva and Barton, 2018). Finally, the sweep of Neanderthal alleles up in frequency may not be due to a beneficial Neanderthal allele, but instead hitch-hiking with a new beneficial allele that arose via mutation more recently than the Neanderthal introgression, by chance on the background of a Neanderthal haplotype. If Neanderthal-introgressed alleles are at ~ 2% frequency genome-wide, then we should expect about 2% of recent sweeps from new mutation to arise on a Neanderthal-introgressed background and sweep them to high frequency.

We identify two regions as having clear statistical support for very recent selection (OAS1,OAS3 and CACNA1S,ASCL5). These regions have very high 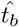, which our simulations show are very rarely estimated under cases of early selection. We are less convinced of alternative explanations for their late rise in frequency because they would need to be replicated in multiple populations. Specifically, given that the Neanderthal alleles rose in frequency so late, and that in both of these regions Neanderthal ancestry is at high frequency in present day Europeans and East Asians, this rise must have begun after Europeans and East Asians diverged from each other, i.e. independently. The chance that the Neanderthal alleles both recombined off of their deleterious background or both hitchhiked on a sweep in the region are extremely small. Thus, for OAS1,OAS3 and CACNA1S,ASCL5, we have evidence that Neanderthal alleles were independently beneficial in Europeans and East Asians, well after admixture with Neanderthals.

In the CACNA1S,ASCL5 region, the estimated waiting time until selection 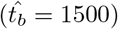 implies selection began after the sampling time of the West Eurasian Upper Paleolithic, however this population does carry Neanderthal haplotypes at high frequency in this region. The Neanderthal allele may have drifted away from low frequency in this population after it diverged from the European ancestry populations that we study, which our simulations show happens reasonably often and does not confound our method. Alternatively, selection may have started earlier in the common ancestor of European ancestry populations and more recently in East Asian ancestry populations. We reran our method after removing East Asian samples but our estimates did not change, which could either reflect a selection onset more recent than the West Eurasian Upper Paleolithic were sampled or poor performance of the method in the absence of East Asian samples. The latter possibility was beyond the scope of this paper to investigate, however future applications could investigate the effect of sampling and extend the method to allow for different onset times when selection independently favors alleles. In total, Neanderthal alleles in this region appear to have been independently favored in the ancestors of East Asians and the ancestors of Europeans, indicating that geographically separated populations experienced similar changes in environmental conditions, possibly at different times.

The selection pressure favoring the OAS1,OAS3 haplotype is thought to be related to innate immunity. Indeed, a recent study found that the Neanderthal haplotype at OAS1 is protective against COVID-19 severity, hospitalization, and susceptibility (Zhou et al., 2020). It has previously been suggested that this gene contributes to traits under balancing selection because haplotypes at this locus have shown variable expression responses to different Flaviviruses (Sams et al., 2016) and a separate Denisovan haplotype segregates at high frequency at this locus in Melanesians (Mendez et al., 2012). Our recent date of selection, and the fact that genomic samples dated closer to the admixture time do not carry Neanderthal haplotypes in this region, are consistent with a recent onset of positive selection in a longterm cycle of balancing selection. A number of other genes known to influence immunity such as the TLR6-TLR1-TLR10 cluster and HLA class I genes harbor candidates of adaptive introgression, however we did not analyze them as they contain both Neanderthal and Denisovan haplotypes in Eurasian populations (Abi-Rached et al., 2011; Dannemann et al., 2016). The timing of adaptation for the OAS1,OAS3 region is at odds with the hypothesis that the transfer of pathogens from Neanderthals to modern humans necessitated rapid human adaptation using Neanderthal-derived immunity alleles (Enard and Petrov, 2018; Greenbaum et al., 2019). However, it would be interesting to test this idea more generally by attempting to date the start of adaptation for more of these introgressed, putative immunity-related haplotypes.

Our investigation contributes to a growing model-based understanding of the genomic patterns of adaptive introgression. Setter et al. (2020) introduced VolcanoFinder, a method that identifies cases of adaptive introgression by probing the recipient population’s patterns of heterozygosity around the selected site. Using expected coalescence times and assuming fixation of the selected allele, they predict very low diversity in a small window around the selected site and very high levels of diversity at intermediate distances. Our predicted within-population probabilities of coalescing characterize similar patterns when combining results from both partitions of a selected population (weighted by their frequency in the population, i.e. the frequency of the selected allele). However, the characteristic volcano pattern would disappear as the frequency of the selected allele decreases or waiting time until selection increases. Our work demonstrates that both of these situations are frequent, and thus the continued difficulty of developing relaxed tests for adaptive introgression while avoiding false positives. Shchur et al. (2020) also use a coalescent approach, in their case to model how adaptive introgression affects the distribution of introgression tract lengths. Fixing the time until introgression and admixture proportion, they show how stronger selection increases the introgression tract lengths. While we do not explicitly model tract lengths, these patterns are apparent in our coalescent predictions.

We implemented a flexible inference framework that could be further modified to investigate the spatio-temporal history of adaptation in modern humans and other systems, whether via introgression or derived mutations. Future applications could distinguish among groups of truly selected populations, allow the frequency of the selected allele to vary among populations, or to distinguish between drift and selection. Our models and estimation method for the timing of adaptation via introgression can be generalized to other organismal systems. In our current application we assume that the introgressed lineage coalesces with the Neanderthal lineage due to the Neanderthal population’s low effective population size, as this allows us to side step modeling the sweep in Neanderthals. However, in other systems the donor population cannot be assumed to have a very low effective population size, but it should be simple to include the extra terms modeling the sweep-induced increase in coalescence between the recipient and donor populations close to the selected site. The application of methods like this to other species would allow a more general understanding of how often introgression is rapidly favored versus supplying genetic variation for future adaptation.

Using a combination of ancient DNA and haplotype based timing, we documented spatially and temporally varying selection on Neanderthal alleles. Our results provide evidence that admixture between diverged populations can be a source of genetic variation for adaptation in the long term. They also allow us to better understand the historical and geographic contexts within which selection favored Neanderthal introgression. As experimental work begins to identify the specific alleles and phenotypes potentially selected on, we can more fully flesh out how interbreeding with archaic populations tens of thousands of years ago has shaped the evolution of modern human populations.

## Acknowledgements

We thank Iain Mathieson for helpful feedback and for sharing genotype likelihoods from the ancient samples that we included in our analysis. We also thank members of the Coop lab for helpful discussions and Erin Calfee, Chuck Langley, Pavitra Muralidhar, Matthew Osmond, Anita To, Michael Turelli, and Carl Veller for comments on earlier drafts of our manuscript. We thank Nicholas Barton, Rasmus Nielsen, and three anonymous reviewers for valuable feedback on earlier drafts as well. This work used the Extreme Science and Engineering Discovery Environment (XSEDE) computing resource PSC Bridges through allocation TG-MCB200074 to SY. This work was supported by the National Science Foundation (IOS-1353380 and PGR-1546719) and the National Institutes of Health (GM108779 and GM136290 to GC and GM121372 to Molly Przeworski).

## A Appendix

### A.1 Simulation Details

#### A.1.1 Selection Simulations

Keeping demography constant, we simulated different combinations of *t_b_, s, t_s_*, and set of selected populations (Table S5 and Table S6). After the divergence between the ancestors of Neanderthals and modern humans, we added a mutation with s = 0.025 to a single Neanderthal haplotype at the selected site. It became neutral before admixture with modern humans. We restarted any runs in which the selected mutation did not reach fixation in the Neanderthal population prior to admixture, was not segregating in all selected populations at the onset of selection, was lost from any selected population during the sweep, was not at greater than 20% frequency in at least one selected population at the sweep finish, or was not at greater than 20% frequency in either the European or East Asian population sampled in the present day. The 20% frequency cut off was motivated by the same criteria with which adaptive introgression candidates were identified (Racimo et al., 2017).

As frequency trajectories in SLiM are stochastic, we chose combinations of *s* and ts that would target different final frequencies (*x_s_*) under a deterministic trajectory. For each s and target *x_s_*, we calculated *t_s_* according to equation 4 and assuming that the frequency at the sweep start is the admixture proportion we used in simulations (*g* = 0.02). However, because our simulations condition on the selected mutation segregating in all selected populations at the selection onset, the average starting frequency in our simulations tended to be greater than g, and increased the later selection began. Therefore, our chosen ts on average led to rather high *x_s_* in cases of late selection, even when we targeted *x_s_* = 0.2. Alternative approaches to target a certain low *x_s_* in our simulations would have led to many simulations where a target *x_s_* was reached due to genetic drift rather than natural selection. Those approaches are inappropriate for us, as we focus on understanding the history of natural selection, rather than distinguishing between genetic drift and natural selection.

Of all polymorphic sites that were not filtered, we randomly down-sampled the data set to 12,000 sites, which is close to the number of sites analyzed for data in our application with loci of the same size. As described in the main text, we used a different sampling approach for simulations generated to identify the CLR_*t*_*b*_>0_ boundary for the regions we analyzed in our application.

#### A.1.2 Neutral Simulations

We generated simulations under neutrality using the same demographic history as the selection simulations so that we could estimate the “genome-wide” neutral **F**. We simulated 2000 independent trees by separately recording the tree sequences of 1 bp loci and subsequently overlaying multiple mutations. From these 2000 independent “loci” we estimated the neutral **F** from 19,216 sites after filtering.

### A.2 Estimating probabilities of coalescing from allele frequencies

Using allele frequencies, we obtain unbiased estimates of **F**, our genome-wide neutral probabilities of coalescing. Rearranging the equation providing the covariance of the change in population allele frequencies from the ancestral population, the probability a pair of ancestral lineages coalesce before the root is

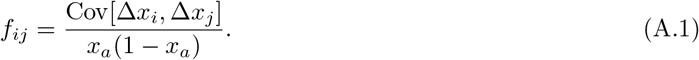

Since Δ*x_i_* = *x_i_* − *x_a_*, the above covariance equals 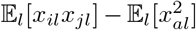. In our scripts we estimate *f_ij_* using an equivalent expression that directly averages over the two possible alleles at a locus,

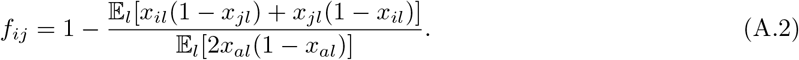

The numerator of the fraction represents the probability of sampling different alleles from population *i* and population *j*, which we estimate by taking its average over all loci. When *i* and *j* correspond to the populations that root the tree, this average serves as the unbiased estimator of the denominator, 2*x_a_*(1 − *x_a_*). When *i* = *j*, such that the numerator represents the expected heterozygosity within a population, we must account for how allele frequencies change when we sample without replacement, i.e. we must remove the finite sampling bias. If *n_il_* represents the sample size of population *i* at locus *l*, then when *i = j*,

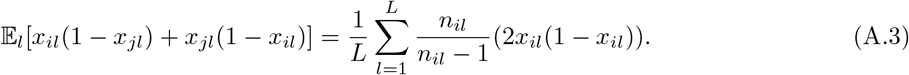

When validating that our model predictions match the probabilities of coalescing in our selection simulations, such as in Figure 4, we repeated the above procedure for sets of loci binned by genetic distance to the selected site.

### A.3 Inference Details

When we calculate likelihoods of population allele frequencies, we need to account for the finite sampling bias in our estimates. Previously, we described the variance in the true population allele frequency due to genetic drift, however there is an additional variance in our estimates due to sampling. Since the count of a certain allele type in a sample is binomially distributed according to the true allele frequency, then with sample size *n_il_* the variance in allele frequency due to sampling is expected to be approximately 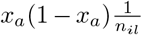. Thus the total variance in the population allele frequency is 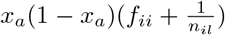. Therefore, when we calculate the probability of population allele frequencies at a locus, we first modify **F**^(**S**)^ by adding 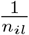 along the diagonal for each population *i*.

Since we do not know the ancestral allele frequency at each locus, we approximate it with the mean across our sampled population allele frequencies,

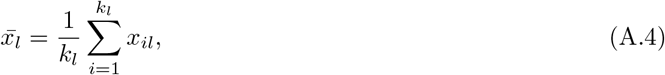

where *k_i_* is the number of populations with allele frequency data at locus l. We acknowledge that in our case 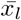 will be biased toward European allele frequencies. With allele frequencies now distributed relative to each other, we accordingly modify the population allele frequencies and covariance matrix by mean-centering them. The new population allele frequencies therefore represent deviations from their mean, and a negative covariance among pairs of populations implies that we predict their allele frequencies to be on opposite sides of this mean. As 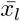 uses information from all populations, we lose a degree of freedom and thus drop information from a single population. Our resulting mean-centered allele frequencies are

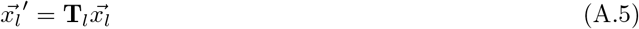

and our mean-centered covariance matrix is

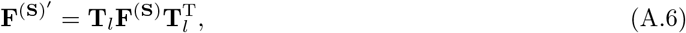

where **T**_*l*_ is the *k_l_* − 1 by *k_l_* matrix with 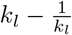 on the diagonal and 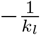 elsewhere. Therefore we calculate the probability of mean-centered allele frequencies at a locus as

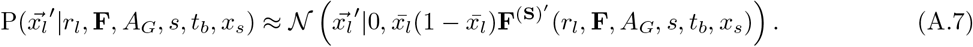

We take the same approach when calculating probabilities under the null model.

To allow for computational efficiency, we first bin each site’s absolute genetic distances to the selected site into 10^−3^ cM intervals. Each bin’s midpoint value is then used as the representative recombination rate to predict **F**^(**N**)^ and **F**^(**S**)^ for each parameter combination. When we calculate the probability of allele frequencies at each site, we use its bin’s representative **F** and remove the rows and columns corresponding to unsampled populations at the site prior to sample size correction and mean centering.

## S1 Supplementary Tables

Table S1: Not shown here. Detailed information on ancient samples are included in a supplementary CSV file.

**Table S2:** Timing and proportion of admixture between listed ancient populations and CEU (a European-ancestry population). Admixture proportions were determined from Haak et al. (2015). Simulations use adjusted admixture proportions so that CEU has the listed admixture proportions following the final admixture event. Our models assume a single time for all admixture among European-ancestry populations, which we define as the average of the migration times listed here.

| Population | Migration Time<br>(generations<br>ago) | Admixture Pro-<br>portion |
| --- | --- | --- |
| EF | 224 | 0.52 |
| Steppe | 155 | 0.36 |
| WHG | 226 | 0.12 |

**Table S3:** Putative regions of adaptive introgression that we analyzed. Genes correspond to those listed with signals of adaptive introgression in Racimo et al. (2017). We label intergenic regions by their chromosome and midpoint position.

| Chrom. | Start to End Positions<br>(hg19) | Genes | Plot Label |
| --- | --- | --- | --- |
| 1 | 193772128 - 193999736 |  | chr1:193885932 |
| 1 | 195854615 - 195922174 |  | chr1:195888394 |
| 1 | 201038832 - 201102553 | CACNA1S, ASCL5 | CACNA1S,ASCL5 |
| 2 | 65395511 - 65533682 | ACTR2 | ACTR2 |
| 2 | 68347097 - 68507952 | PPP3R1, PNO1 | PPP3R1,PNO1 |
| 2 | 119536920 - 119693701 | EN1 | EN1 |
| 2 | 154435207 - 154551882 |  | chr2:154493544 |
| 2 | 159794421 - 160204173 | TANC1, WDSUB1,<br>BAZ2B | TANC1...BAZ2B |
| 2 | 160896628 - 161078020 | PLA2R1, ITGB6 | PLA2R1,ITGB6 |
| 4 | 5828060 - 5920067 | EVC, CRMP1 | EVC,CRMP1 |
| 4 | 189157618 - 189216507 |  | chr4:189187062 |
| 5 | 507500 - 653905 | SLC9A3 | SLC9A3 |
| 5 | 167971758 - 168166025 | SLIT3 | SLIT3 |
| 6 | 1396838 - 1451472 | FOXF2 | FOXF2 |
| 6 | 66231951 - 66532520 | EYS | EYS |
| 7 | 129861223 - 129929228 | CPA2 | CPA2 |
| 7 | 136570934 - 136667889 | CHRM2 | CHRM2 |
| 8 | 13846885 - 13942904 |  | chr8:13894894 |
| 8 | 13946965 - 14174597 | SGCZ | SGCZ |
| 9 | 16596898 - 16845056 | BNC2 | BNC2 |
| 11 | 11552692 - 11625808 | GALNT18 | GALNT18 |
| 11 | 70891472 - 70976597 |  | chr11:70934034 |
| 12 | 54151324 - 54244563 |  | chr12:54197944 |
| 12 | 84808525 - 84998584 |  | chr12:84903554 |
| 12 | 113337379 - 113427536 | OAS1, OAS3 | OAS1,OAS3 |
| 14 | 100944952 - 101071031 | WDR25, BEGAIN | WDR25,BEGAIN |
| 15 | 43432954 - 44053859 | TGM5, TGM7 | TGM5,TGM7 |
| 15 | 74702840 - 75030779 | SEMA7A, UBL7 | SEMA7A,UBL7 |
| 15 | 85908189 - 86341507 | AKAP13, KLHL25 | AKAP13, KLHL25 |
| 16 | 51306915 - 51398178 |  | chr16:51352546 |
| 16 | 77959647 - 78071863 | VAT1L | VAT1L |
| 18 | 60124891 - 60268585 | ZCCHC2 | ZCCHC2 |
| 18 | 60658804 - 60777888 |  | chr18:60718346 |
| 19 | 33510019 - 33763643 | RHPN2, GPATCH1,<br>WDR88, LRP3,<br>SLC7A10 | RHPN2...SLC7A10 |
| 20 | 62156560 - 62210005 | PPDPF, PTK6, HELZ2 | PPDPF...HELZ2 |
| 22 | 49077496 - 49144645 | FAM19A5 | FAM19A5 |

**Table S4:** Proposed parameters of *s* and *t_b_*, each combination of which we calculated composite likelihoods. For a given run of the method, we did not evaluate a combination of s and *t_b_* if the sweep would not complete (deterministically) before the present day, i.e. if the time until selection is recent enough and/or the sweep time *t_s_* (which also depends on *x_s_*) is long enough such that *t_b_* + *t_s_* > *t_i_*.

| Proposed selection coefficients ($s$) | Proposed waiting times until selection ($t_b$) |
| --- | --- |
| 0.002, 0.005, 0.01, 0.015, 0.02, 0.025, 0.03, 0.05 | 0, 100, 200, 300, 400, 500, 600, 700, 800, 900, 1000, 1100, 1200, 1300, 1400, 1500, 1600, 1700 |

**Table S5:** Simulated combinations of waiting time until selection (*t_b_*), group of selected populations, selection coefficient s, and sweep phase duration (*t_s_*). Combinations of *s* and *t_s_* are labeled by numbers in Table Table S6.

| Waiting time<br>until selection ($t_b$) | Selected human populations | Combinations of $s$ and $t_s$<br>(references rows in Table S6) |
| --- | --- | --- |
| 0 | All Eurasian | 1, 2, 3, 4, 5 |
| 200 | All Eurasian | 7, 8, 9 |
| 400 | All Eurasian | 7, 8, 9 |
|  | All European | 6, 7, 8, 9 |
| 600 | All Eurasian | 7, 8, 9 |
|  | All European | 6, 7, 8, 9 |
|  | All Eurasian except EurUP | 6, 7, 8, 9 |
| 800 | All Eurasian | 7, 8, 9 |
|  | All Eurasian except EurUP | 6, 7, 8, 9 |
|  | All Eurasian except East Asia, EurUP | 6, 7, 8, 9 |
| 1000 | All Eurasian except EurUP | 6, 7, 8, 9 |
|  | All Eurasian except East Asia, EurUP | 6, 7, 8, 9 |
| 1200 | All Eurasian except EurUP | 6, 7, 8, 9 |
|  | All Eurasian except East Asia, EurUP | 6, 7, 8, 9 |
| 1400 | All Eurasian except EurUP | 6, 7, 8, 9 |
|  | All Eurasian except East Asia, EurUP | 6, 7, 8, 9 |

**Table S6:** Combinations of *s* and *t_s_* used in selection simulations used to evaluate the method’s performance. Target *x_s_* corresponds to the average frequency the selected allele would reach at the end of the sweep phase if it started the sweep at frequency *g* = 0.02. The last column (Runs) provides the number of replicates simulated for a provided waiting time until selection and group of populations experiencing selection.

| | Selection Coefficient ($s$) | Sweep Phase Duration ($t_s$) | Target $x_s$ | Runs |
| --- | --- | --- | --- | --- |
| 1 | 0.005 | 501 | 0.20 | 700 |
| 2 | 0.005 | 697 | 0.40 | 700 |
| 3 | 0.010 | 349 | 0.40 | 700 |
| 4 | 0.010 | 430 | 0.60 | 700 |
| 5 | 0.010 | 528 | 0.80 | 700 |
| 6 | 0.005 | 501 | 0.20 | 100 |
| 7 | 0.005 | 655 | 0.35 | 200 |
| 8 | 0.010 | 327 | 0.35 | 200 |
| 9 | 0.010 | 474 | 0.70 | 100 |

## S2 Method Details

### S2.1 Method Validation

The method’s power to reject immediate selection increases with the true waiting time until selection (*t_b_*) and the average frequency of the selected allele among putative selected populations (*x_s_*) because they each lead to later estimates of *t_b_*, which correspond to higher CLR_*t*_*b*_>0_ (Figures S1–S5). Among the simulations in which our method rejects immediate selection, it reliably identifies late selection when *t_b_* > 800. Notably, it can tell that selection is recent despite the selected allele having drifted to high frequency earlier in time. For example, the West Eurasian Upper Paleolithic (EurUP) were sampled 908 generations after admixture, such that whenever *t_b_* > 900, allele frequencies in that population can only reach higher frequencies because of genetic drift. Despite these high frequency cases when the true *t_b_* > 900, the method can tell that selection started after the EurUP were sampled.

When East Asians are the only Eurasian population that does not carry the allele at high frequency, the method is biased toward inferring earlier selection when *t_b_* < 800 and does poorly when *t_b_* = 800. When EurUP is the only population that does not carry the allele at high frequency, the method is biased toward inferring later selection when *t_b_* < 800.

**Figure S1:**
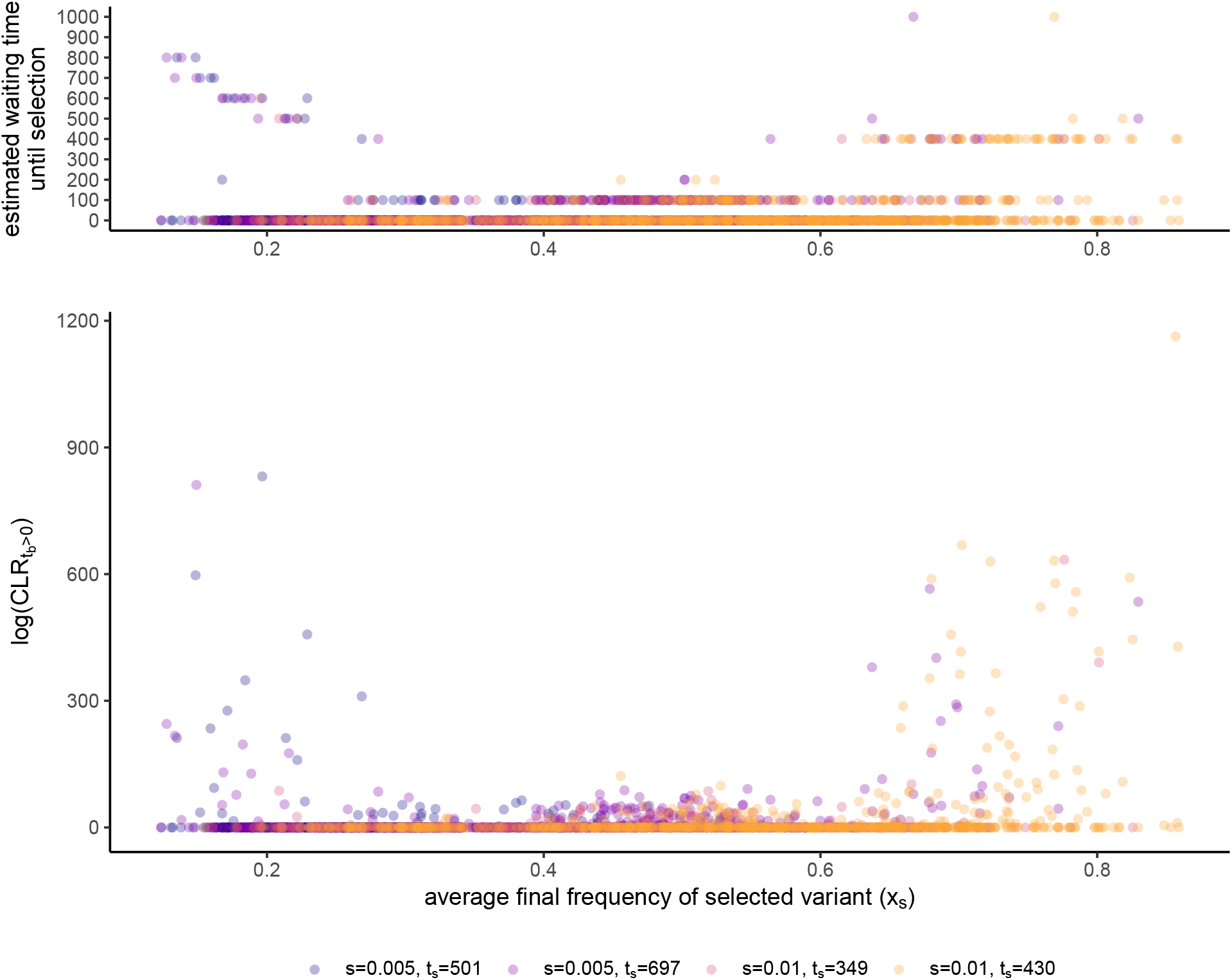
(Top) Relationship between the average final frequency of the selected variant (*x_s_*) and the estimated waiting time until selection 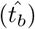 among immediate selection simulations. The method estimated immediate selection 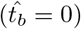 in 85% of these simulations. (Bottom) Relationship between *x_s_* and the log of the composite likelihood ratio between non-immediate and immediate selection, CLR_*t*_*b*_>0_, among immediate selection simulations. logCLR_*t*_*b*_>0_ equals zero for the vast majority of low to intermediate *x_s_* because at these lower frequencies the method is more biased toward inferring immediate selection. The relationship between *x_s_* and logCLR_*t*_*b*_>0_ does not differ among combinations of s and *t_s_*. See Table S6 for targeted *x_s_* corresponding to each combination of *s* and *t_s_*.

**Figure S2:**
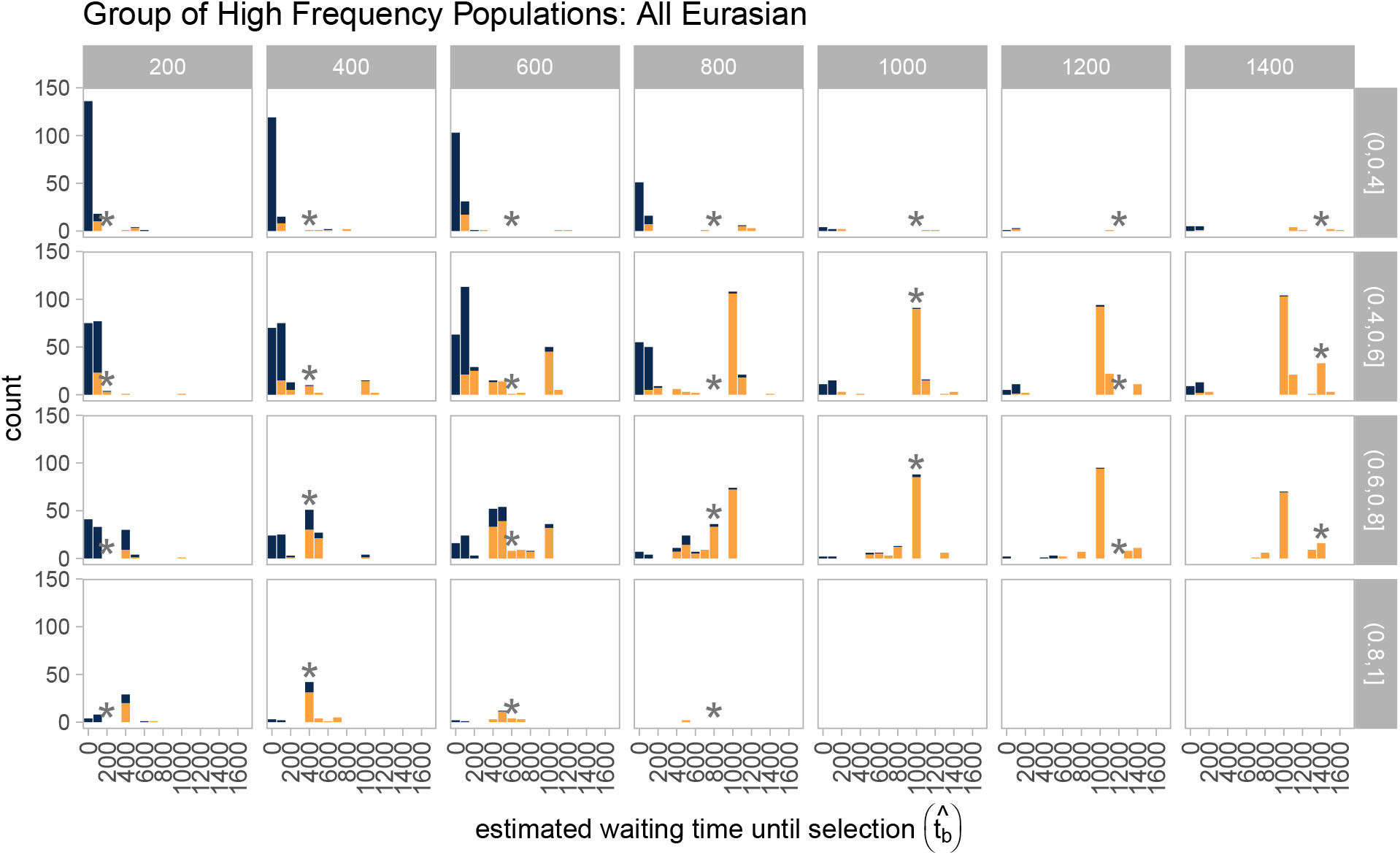
Distribution of estimated waiting times until selection among simulations in which the method considers all Eurasian populations to be selected. Blue bars represent cases in which the method did not reject immediate selection, and orange bars represent cases in which the method did reject immediate selection. Blue bars are stacked on top of orange bars (they do not overlap). Results are shown for each combination of true waiting times until selection (columns) and binned average final frequencies of the selected variant (*x_s_*, rows). Asterisks mark the estimated waiting time until selection that matches the truth.

**Figure S3:**
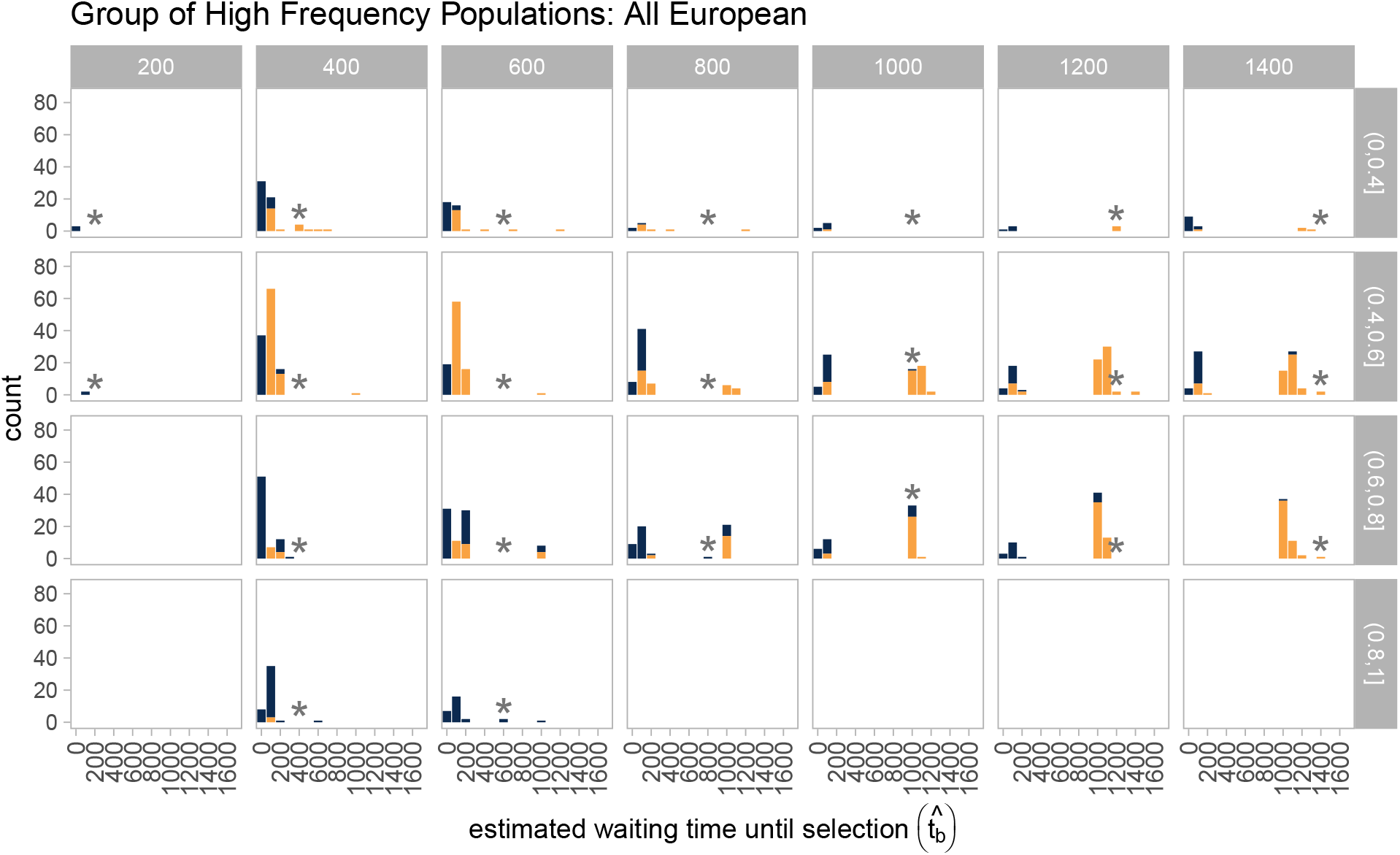
Distribution of estimated waiting times until selection among simulations in which the method considers all European populations (including the West Eurasian Upper Paleolithic) to be selected. Blue bars represent cases in which the method did not reject immediate selection, and orange bars represent cases in which the method did reject immediate selection. Blue bars are stacked on top of orange bars (they do not overlap). Results are shown for each combination of true waiting times until selection (columns) and binned average final frequencies of the selected variant (*x_s_*, rows). Asterisks mark the estimated waiting time until selection that matches the truth.

**Figure S4:**
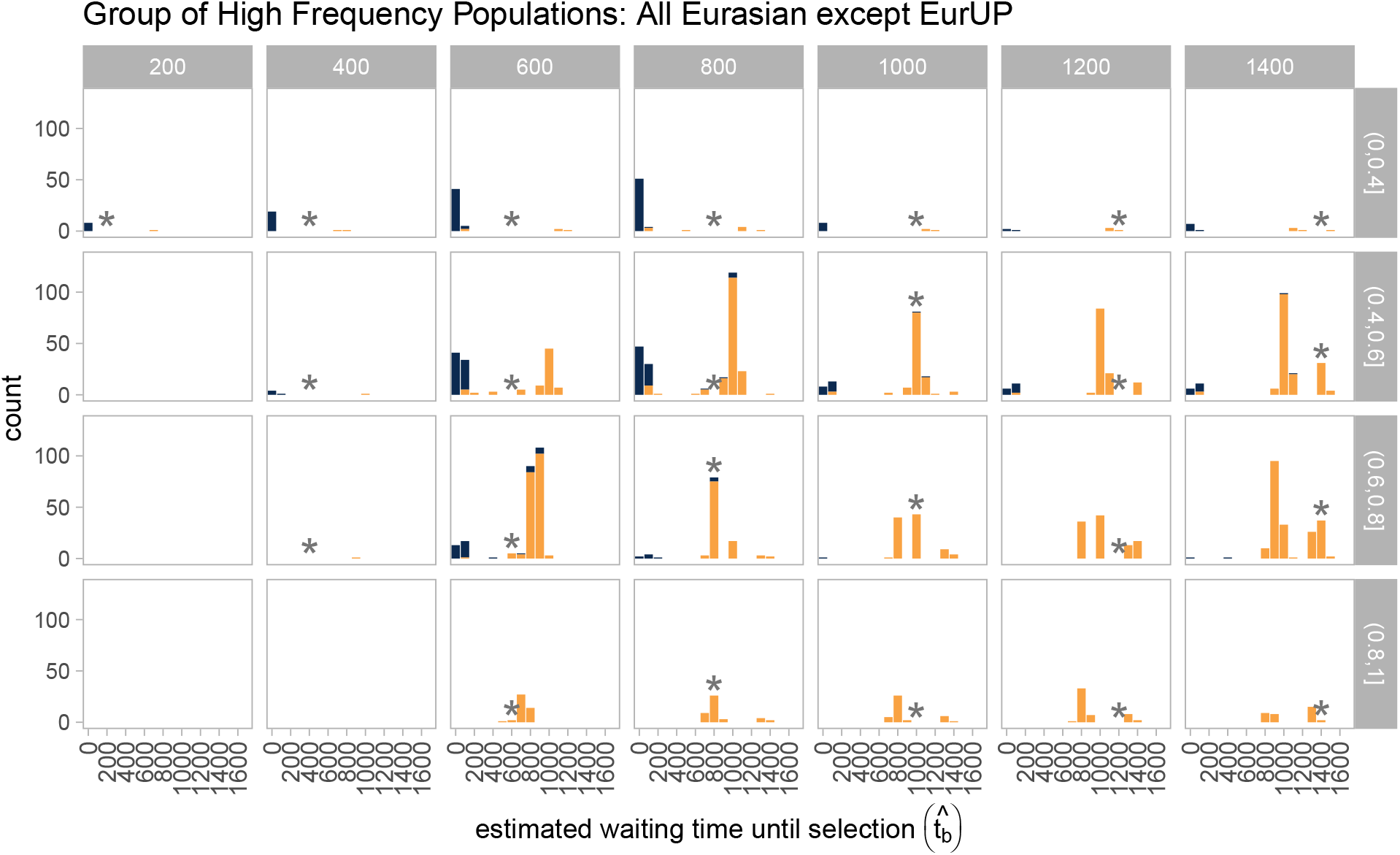
Distribution of estimated waiting times until selection among simulations in which the method considers all Eurasian populations except the West Eurasian Upper Paleolithic to be selected. Blue bars represent cases in which the method did not reject immediate selection, and orange bars represent cases in which the method did reject immediate selection. Blue bars are stacked on top of orange bars (they do not overlap). Results are shown for each combination of true waiting times until selection (columns) and binned average final frequencies of the selected variant (*x_s_*, rows). Asterisks mark the estimated waiting time until selection that matches the truth.

**Figure S5:**
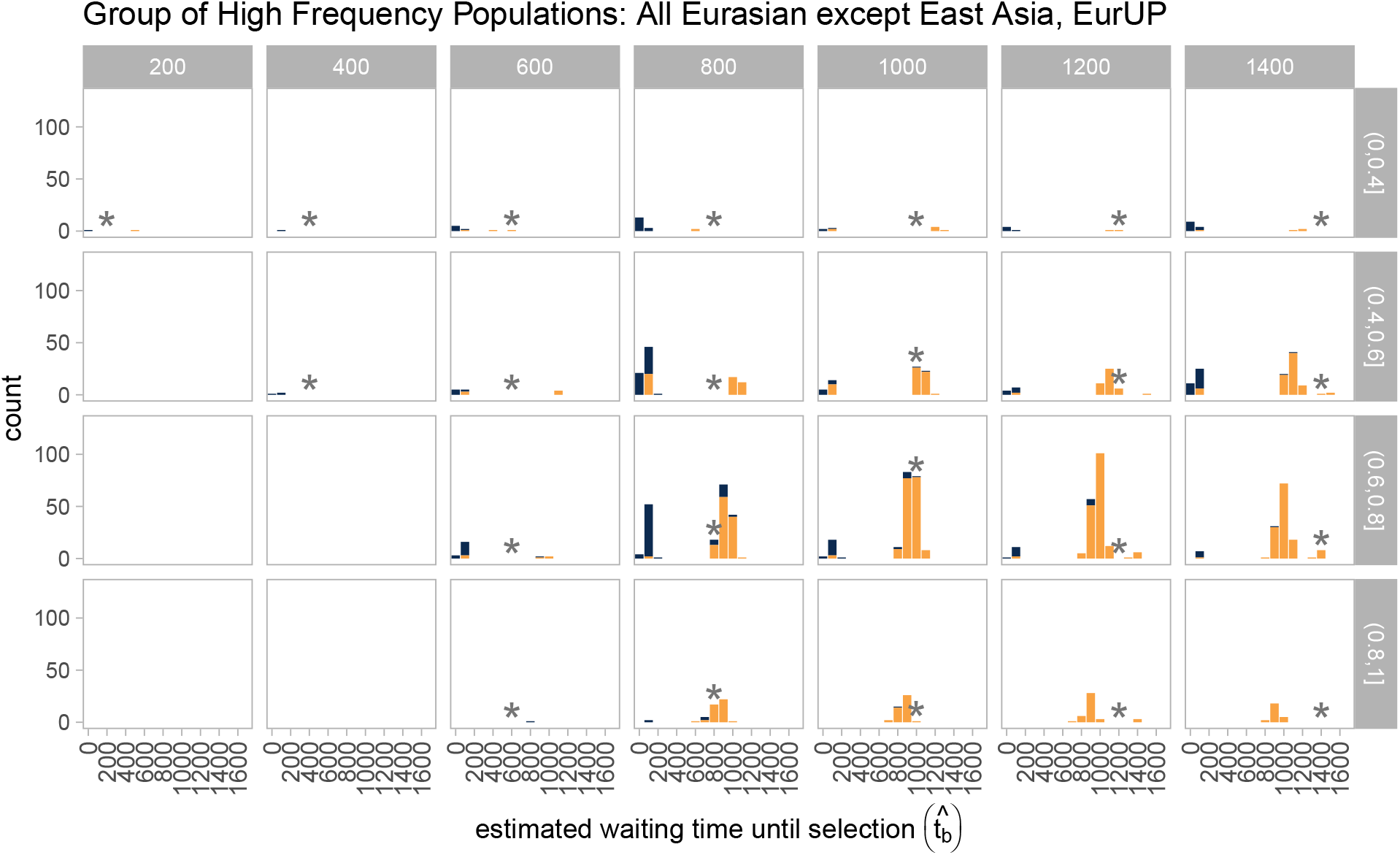
Distribution of estimated waiting times until selection among simulations in which the method considers all Eurasian populations except the West Eurasian Upper Paleolithic and East Asians (represented by CHB) to be selected. Blue bars represent cases in which the method did not reject immediate selection, and orange bars represent cases in which the method did reject immediate selection. Blue bars are stacked on top of orange bars (they do not overlap). Results are shown for each combination of true waiting times until selection (columns) and binned average final frequencies of the selected variant (*x_s_*, rows). Asterisks mark the estimated waiting time until selection that matches the truth.

**Figure S6:**
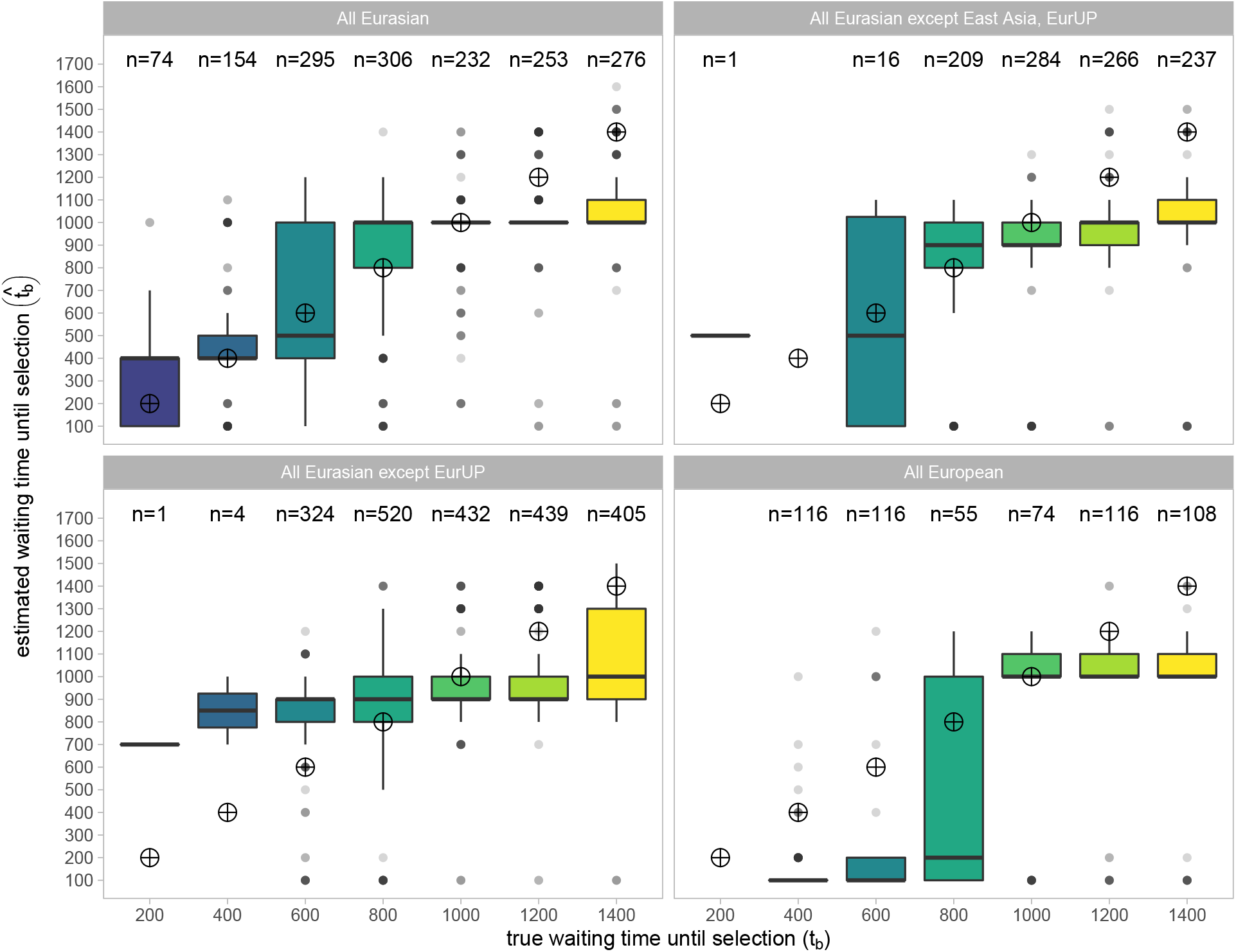
Estimated waiting time until selection 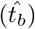 among simulations in which the method rejected immediate selection. Target points mark the true waiting time until selection that we aim to estimate. Earlier true waiting times have fewer observations because the method has less power to reject immediate selection in this parameter space. Each panel corresponds to a different group of populations that the method considers selected.

**Figure S7:**
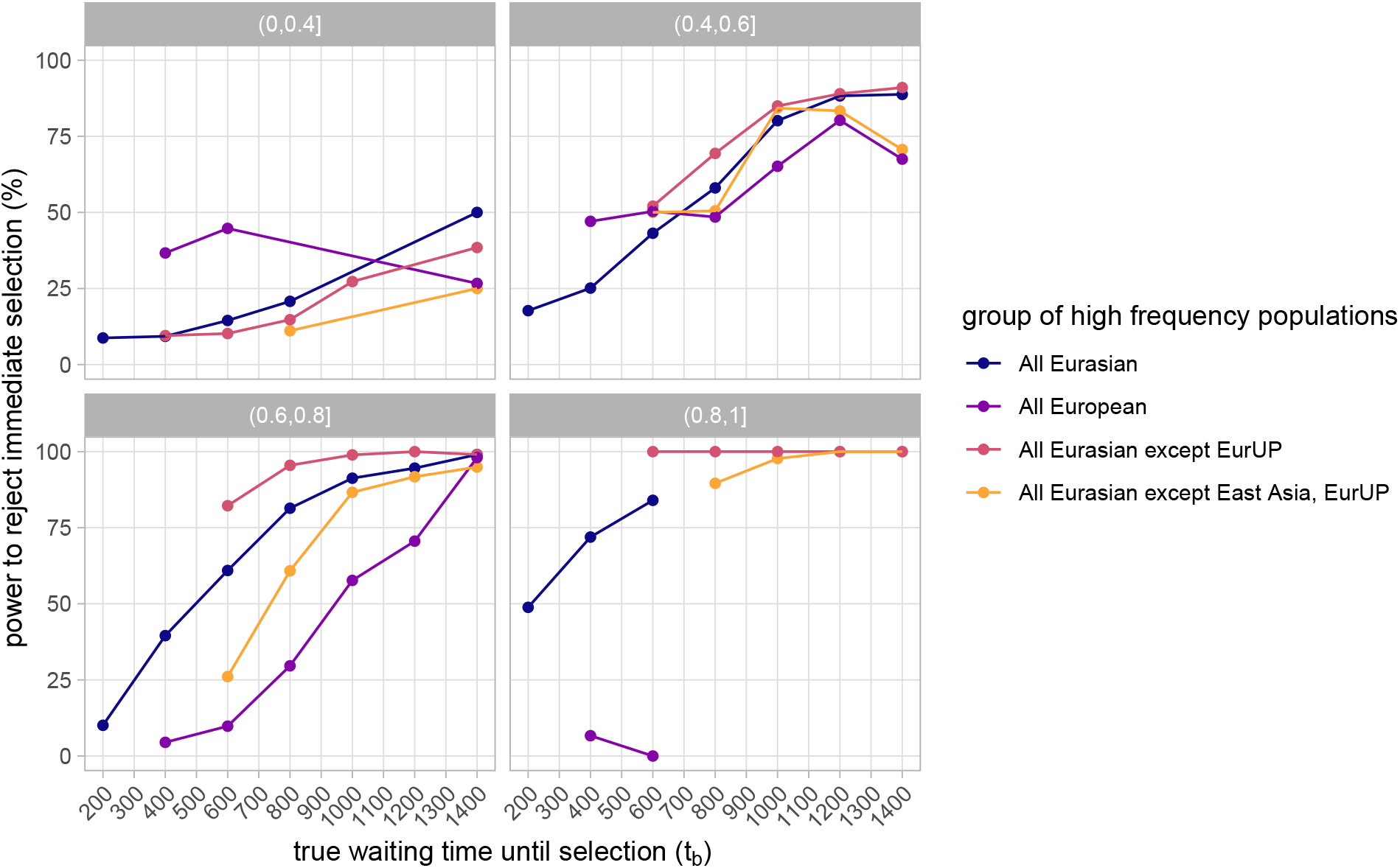
Power to reject immediate selection as the true waiting time until selection (*t_b_*) increases. Lines correspond to simulations with different groups of populations considered selected. Points with fewer than 10 observations were removed. Panels correspond to bins of *x_s_*, the selected allele’s average frequency among populations considered selected.

**Figure S8:**
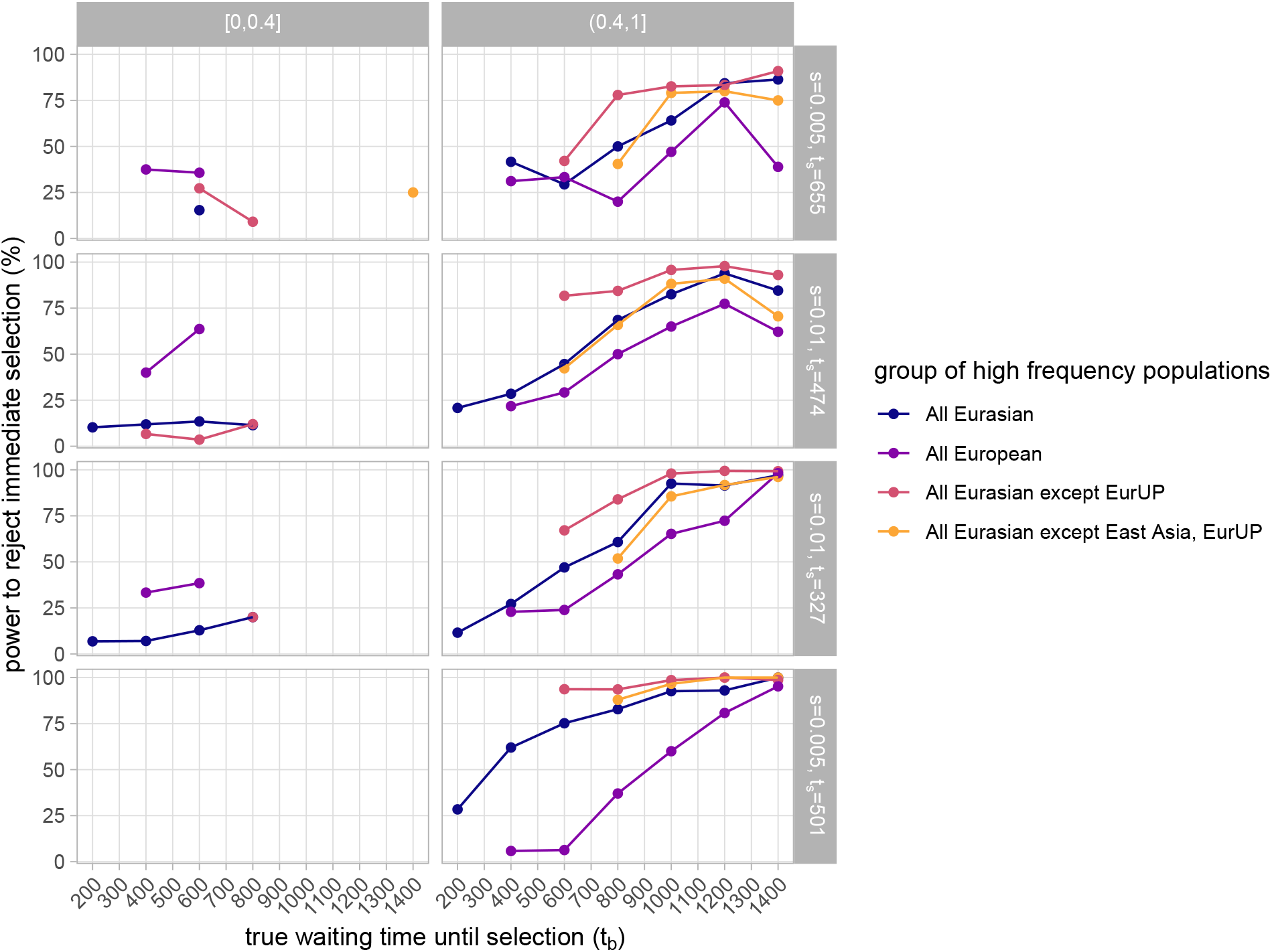
Power to reject immediate selection as the true waiting time until selection (*t_b_*) increases. Lines correspond to simulations with different groups of populations considered selected. Points with fewer than 10 observations were removed. Columns correspond to bins of *x_s_*, the selected allele’s average frequency among populations considered selected, and rows correspond to combinations of s and ts used in simulations.

**Figure S9:**
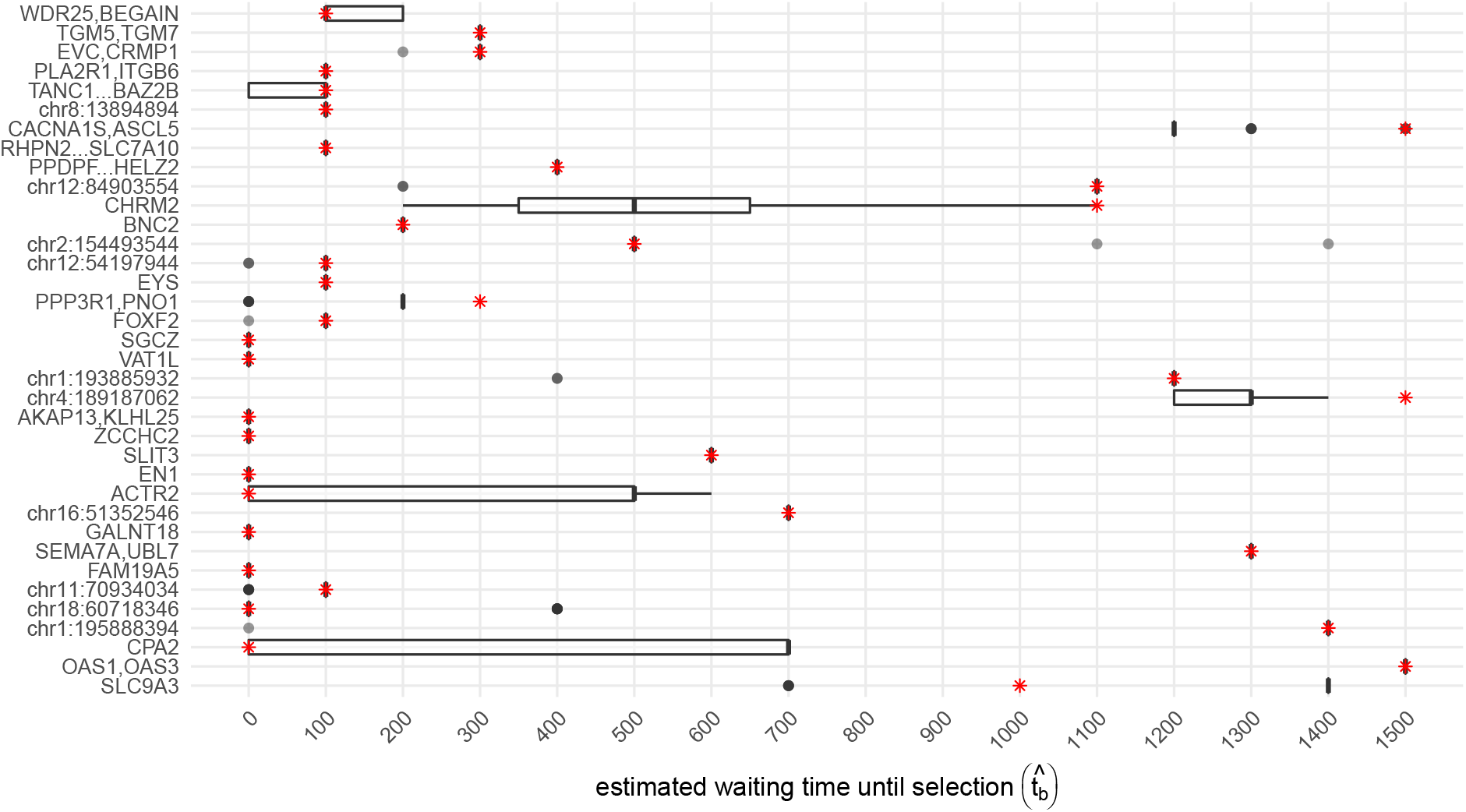
Each region’s estimated waiting time until selection 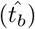 when different sets of ancient samples were randomly assigned to each population’s genotype partitions according to their posterior genotype probabilities from imputation. Boxplots show the distribution under 40 resampled datasets. The red asterisks mark each region’s 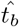 under the dataset in which ancient partitions are assigned based on the maximum likelihood genotypes. Regions are presented in the same order as in Figure 2.

### S2.2 Results under different datasets and demographic assumptions

#### S2.2.1 Inclusion of the Altai Neanderthal Sample

Population structure among Neanderthals and among Neanderthal-introgressed haplotypes could lead to erroneous results if we do not include the true introgressing populations in our analysis. To investigate our method’s sensitivity to our choice of Neanderthal population, we compared our results between using only the Vindija sample and using both the Vindija and Altai samples to calculate allele frequencies for the Neanderthal population. We made a single comparison for each region at the ‘best’ partition site whose maximum composite likelihood under the selection model was greatest relative to the null model. In the vast majority of regions, our estimated waiting times until selection 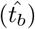 did not change, and if they did they only moved within 200 generations (Figure S10). Our estimates changed greatly in two regions, CHRM2 and chr11:70934034, both of which we have identified elsewhere as difficult to infer their timing of selection onset (see the above and below sections). Our results are likely not sensitive to specification of the Neanderthal population because of their low effective population sizes leading to high probabilities of coalescing before the common ancestor of Neanderthals and modern humans. When the Neanderthal population is composed of Vindija alone, we estimate that the probability of coalescing *f_nn_* = 0.95. For a population composed of Altai and Vindija (n=4), we estimate *f_nn_* = 0.91. Therefore, the ancestral lineages of an allele sampled from the Altai Neanderthal and an allele sampled from the Vindija Neanderthal still have an extremely high probability of coalescing. Since in our models we mainly track whether sampled lineages descended from Neanderthals, such that those lineages coalesce while segregating in a Neanderthal population, the Neanderthal source population has a negligible effect on our predictions.

**Figure S10:**
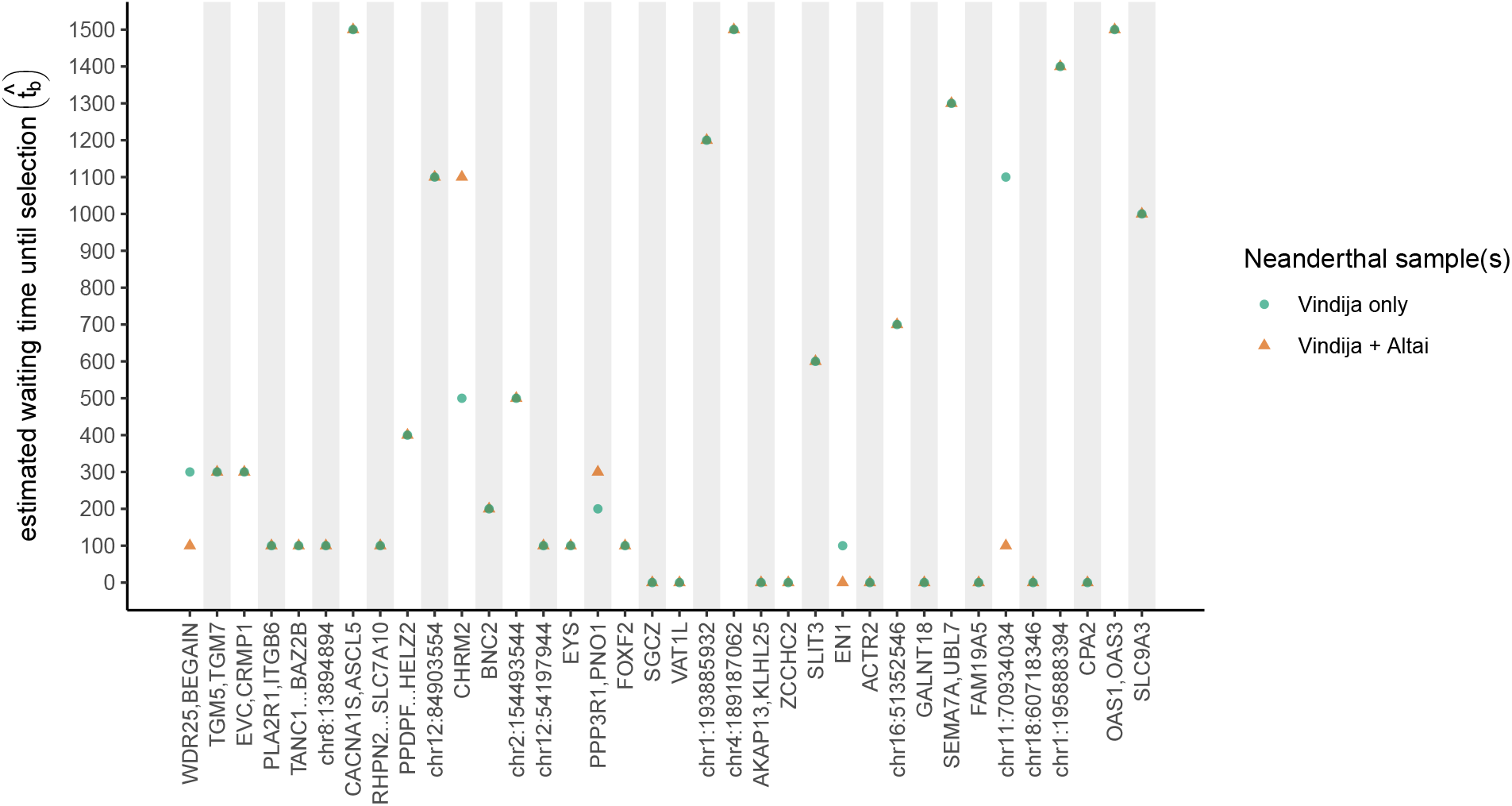
Each region’s estimated waiting time until selection 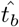 under different sets of samples comprising the Neanderthal population. Regions are presented in the same order as in Figure 2.

#### S2.2.2 Sensitivity to Admixture Graphs

In our application, we ran our method on all partition sites while assuming the admixture graph specified in Figure 1, which we call option A. To evaluate the sensitivity of our estimates to these assumptions, we re-ran the method using three other graphs and the dataset of the highest maximum composite likelihood partition site relative to the null model under option A. Each of these additional graphs contain a single modification from the original. In option B, we changed the time of admixture with Neanderthals from 2068 generations in the past (60kya with a generation time of 29 years) to 1988 generations (55kya with a generation time of 29 years). In option C, we did not model present day Europeans as a mixture of the Bronze Age Steppe, Early Neolithic Farmers, and Western Hunter Gatherers. While there is strong evidence that present day Europeans are indeed a mixture of these three populations, we chose an extreme option to generally demonstrate the insensitivity of our method to specifications of admixture among human populations. In option D, we modified the divergence time among present day Europeans, the Bronze Age Steppe, Early Neolithic Farmers, and Western Hunter Gatherers to be 333 generations (10k years with a generation time of 29 years) later (from 40kya to 30kya). Each region’s timing estimates when assuming each admixture graph are shown in Figure S11.

Overall, estimated waiting times until selection for the majority of regions did not change with each admixture graph, or changed very little. In particular, our modification of the divergence time among European populations (option D) did not change the estimated waiting time until selection from option A with the exception of 3 regions (PPPDF..HELZ2; PPP3R1,PNO1; and chr11:7093403). Generally, estimates more often changed when the admixture time with Neanderthals shifted to be more recent. Among those regions in which there was a change in the estimate from option A when using option C, the estimated waiting time until selection was one or two hundred generations shorter, therefore corresponding to a very similar fixed time that selection began. Thus, changing the admixture time resulted in consistency either in the waiting time until selection or how recently selection began. However, three regions showed large inconsistencies in their estimates as we changed the admixture time, transitioning from very early to very late selection: chr12:54197944, FAM19A5, and chr16:51352546. In the former two regions, we did not reject immediate selection under option A. In chr12:54197944, we would potentially reject immediate selection if we simulated immediate selection under the new admixture graph, and therefore caution that the history of selection in this region is unclear. As for FAM19A5, the composite likelihood ratio between non-immediate and immediate selection remains low such that we would likely still reject immediate selection. In chr16:51352546, we previously rejected immediate selection and likely still would, however we caution that our estimated waiting time until selection is not informative in this region. When we remove any migration among modern human populations, the regions with a large change (> 300 generations) are chr12:84903554, chr1:193885932, ACTR2, SEMA7A,UBL7, and CPA2. Based on the composite likelihood ratio between non-immediate and immediate selection, we would likely continue to reject immediate selection in chr12:84903554, chr1:193885932, and SEMA7A,UBL7. In each of those regions the removal of migration is the only change that alters results, and in each case moves our estimates from recent to early selection. These earlier estimates may be unrealistic given that we have strong evidence for migration among European ancestry populations. It is unclear whether we would continue not to reject immediate selection in ACTR2 and CPA2, though we note these two regions also had wide variation in their estimates when we re-partitioned ancient populations (Figure S9), and so we refrain from making any claims about the timing of selection in these regions.

**Figure S11:**
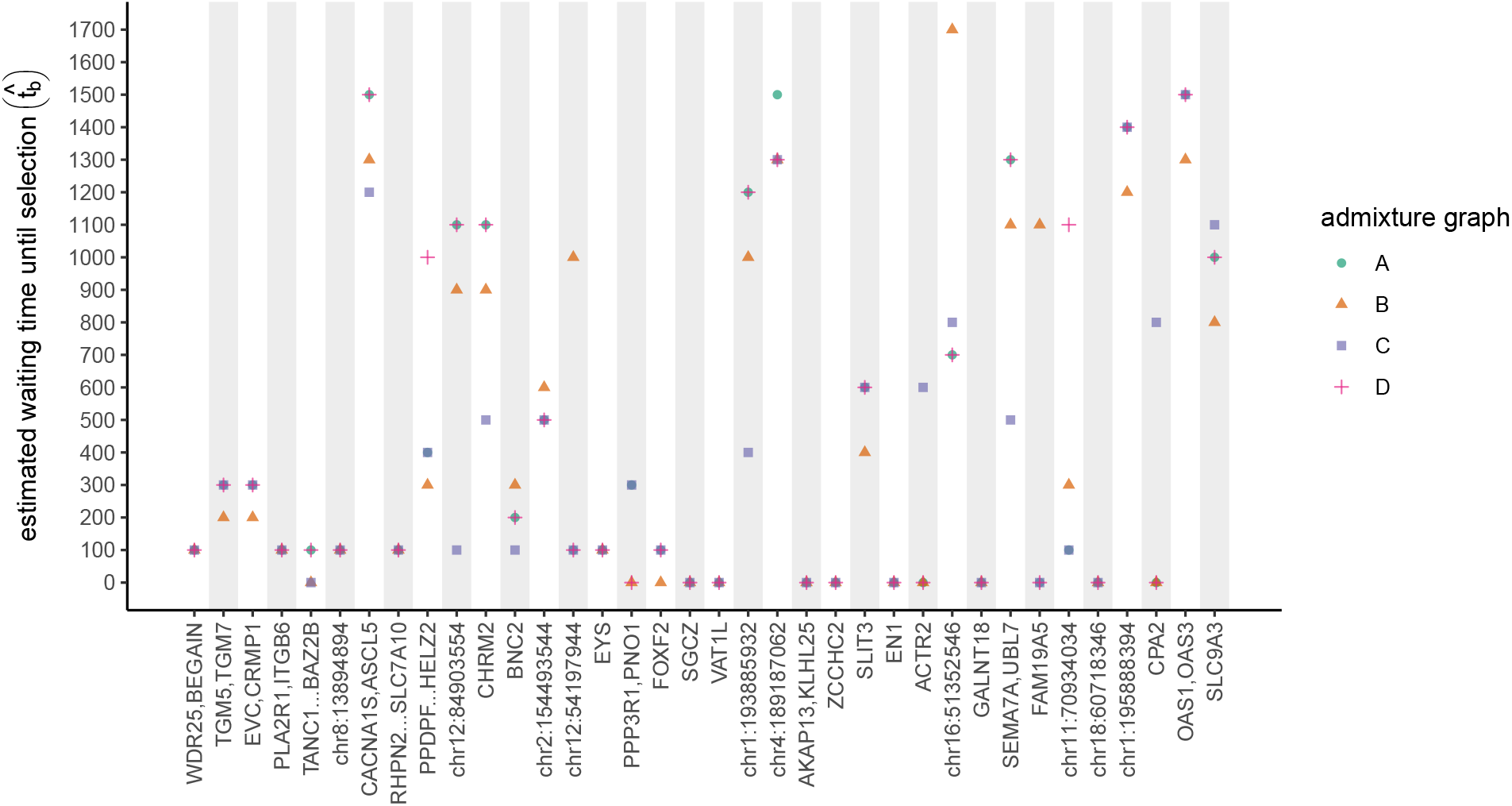
Each region’s estimated waiting time until selection 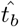 under different admixture graphs. Each admixture graph specified by a letter is introduced in section S2.2.2. Regions are presented in the same order as in Figure 2.

### S3 Model Derivation

#### S3.1 Between selected populations that diverged during neutral phase I

Here, we illustrate another example of how we predict probabilities of coalescing between different categories of populations. We focus in depth on the probability of coalescing between pairs of selected populations who diverged from each other before the sweep started, i.e. during neutral phase I. Afterward, we describe how we change these predictions among other categorical pairs of populations. As a reminder, we consider three phases of the selected allele frequency trajectory: during neutral phase I the selected allele segregates at frequency *g* for *t_b_* generations, during the sweep phase the selected allele rises from frequency *g* to frequency xs over ts generations, and during neutral phase II the selected allele segregates at frequency xs until the present day. Thus neutral phase II lasts a duration of *t_I_* − *t_b_* − *t_s_* generations, where *t_I_* is the time between introgression and the present.

If a pair of populations (*i, j*) diverged from each other *d_ij_* generations in the past, and if this divergence occurred during neutral phase I, then they share a common ancestor for *t_I_* − *d_ij_* generations before the time of introgression. Since the duration of neutral phase I is *t_b_*, we also know that these populations were isolated for the last *t_b_* − *t_I_* + *d_ij_* generations of neutral phase I. To get the probability that the ancestral lineages of an allele sampled in population *i* and an allele sampled in population *j* coalesce, we consider the ancestry background at the selected site that each ancestral lineage is associated with at each phase transition. In the main text, we described the probabilities that a lineage is linked to the selected or non-selected background at the transition between neutral phase II and the sweep completion (Equation 5), in addition to the probability that a lineage remains strictly associated with this ancestry background for the duration of the sweep phase (Equation 6). With probability exp (− (*t_b_* − *t_I_* + *d_j_*)) a lineage does not recombine out of the background that it was associated with at the transition between the sweep start and neutral phase I, until the time at which the two populations share a common ancestor in neutral phase I.

We begin by describing the probability that the pair of lineages coalesce conditional on them both being linked to the selected allele at the transition between neutral phase II and the sweep finish. We say a lineage is “strictly associated” with the sweep if it never recombines out of that background during the sweep phase and does not recombine at all during the preceding generations of neutral phase I when the lineages are in separate populations, such that

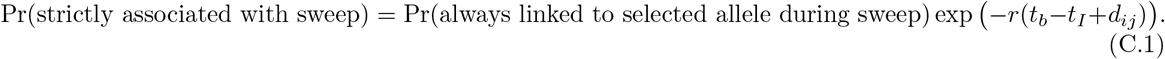

If both lineages remain strictly associated with the sweep, then we consider the possibility that they coalesce because they are associated with a background that is now relatively rare during neutral phase I. This could be due to their descent from the same introgressed allele, or the same non-introgressed allele that became associated with the selected background before the sweep began. As these lineages can only coalesce if they remain on the same background, we treat coalescence and recombination as competing Poisson processes, with recombination occurring at rate 2*r* and coalescence at rate 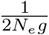, where *N_e_* is the mean effective population size of populations (*i, j*) and *g* is the neutral admixture proportion, or similarly the frequency of the selected background during neutral phase I. The probability that neither of these events occur before the time of introgression is

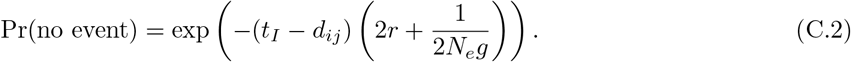

In that case, both lineages are still associated with the selected background at the time of introgression and thus must have descended from Neanderthals, so they coalesce with probability *f_nn_* (the probability a pair of alleles sampled from Neanderthals coalesce). If at least one coalescence or recombination event does occur, we care about the outcome of the first event. The relative probability of coalescence is 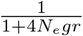 and of recombination is 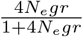. If the event is recombination of one of the lineages, then we condition on whether the other lineage recombines at some point before introgression. With approximate probability exp (−*r*(*t_I_* − *d_ij_*)) it does not recombine, and therefore remains associated with the selected background such that it descended from Neanderthals. Thus the lineages can only coalesce if the lineage that did recombine also descended from Neanderthals, with probability g. If both lineages recombined during this shared portion of neutral phase I, then they coalesce with neutral probability *f_ij_*, which already accounts for whether they are each introgressed or not. We use a similar logic for the remainder of cases that involve one or no lineages remaining strictly associated with the selected background before the pair are in the same population, such that

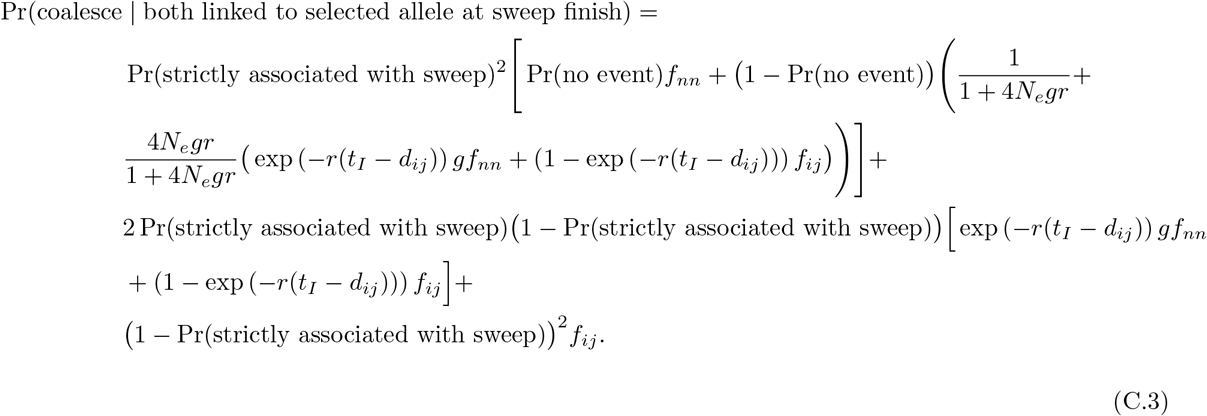

Next we describe the probability that the pair of lineages coalesce conditional on one being associated with the selected allele and the other being associated with the non-selected allele at the transition between neutral phase II and the sweep finish. Since it is relatively rare for the lineage linked to the non-selected allele to switch to the background of the selected allele during the sweep phase, we do not consider the possibility of increased coalescence during neutral phase I. Therefore, we are simply interested in the ancestry background of each lineage at the time of introgression. If neither of them become disassociated with their backgrounds during the sweep phase and do not recombine before the time of introgression, then we know that one allele descended from Neanderthals while the other did not, such that they cannot coalesce. If the lineage originally associated with the non-selected background remains associated with this background during the sweep and does not recombine before introgression, whereas the other lineage does disassociate at some point, they can only coalesce if the disassociating allele did not descend from Neanderthals, with probability 1 − *g*. At this point, we need to slightly modify their probability of coalescing from the neutral estimate because we are dealing with a case in which we exclude descent from Neanderthals as a possibility. The probability that two lineages sampled from Neanderthal admixed populations coalesce, conditional on them both not descending from Neanderthals, is

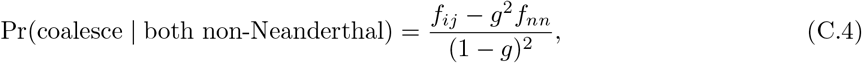

where *g* is the neutral admixture proportion. We have previously described our predictions when one of the lineages descends from Neanderthals due to their consistent association with the selected background while the other lineage may not maintain a consistent association, in addition to when neither lineage maintains a consistent association. Therefore we obtain the following with conditional probabilities ordered in the same manner we discussed them,

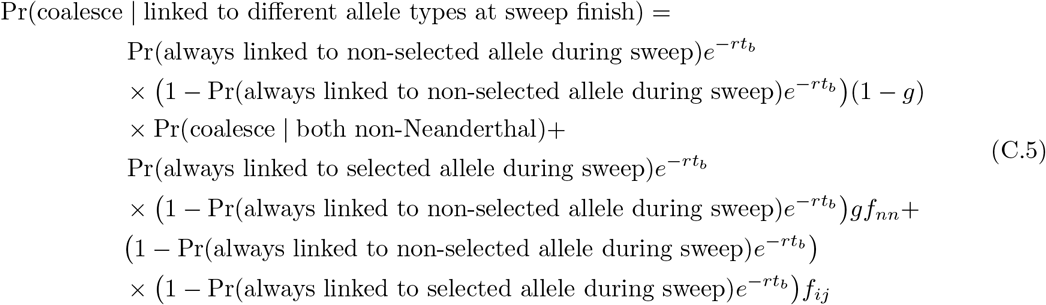

Finally, we describe the probability that the pair of lineages coalesce conditional on both being associated with the non-selected allele at the transition between neutral phase II and the sweep finish. We again condition on their background associations at the time of introgression. If both lineages never disassociate from this background during the sweep and never recombine during neutral phase I, then neither descended from Neanderthals and they coalesce with a slightly higher probability than neutrality as we previously described. If this happens with just one of the lineages, then the other lineage must not have descended from Neanderthals for them to coalesce, again with a slightly higher probability than neutrality. In the remainder of cases, the lineages coalesce neutrally. These cases are summarized in the following,

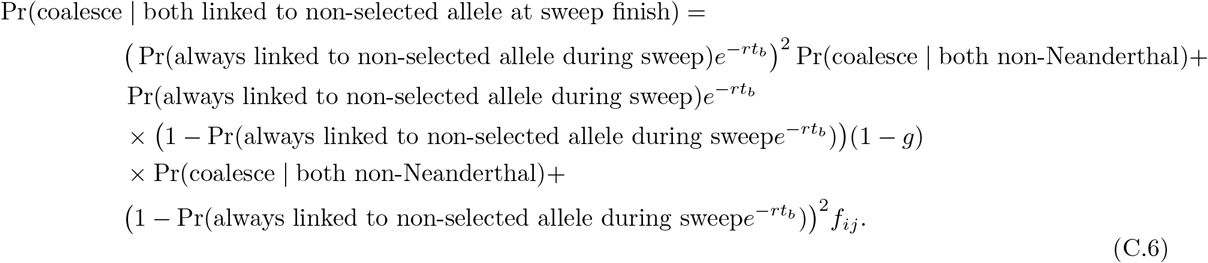

Together, the full probability that a pair of lineages coalesce before the root is

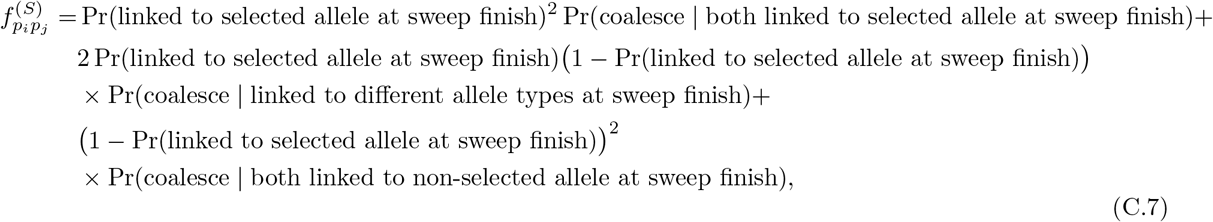

where subscript *p_i_* refers to any partition of population *i* and subscript *pj* refers to any partition of population ^j^.

#### S3.2 Modifications under other scenarios among pairs of selected and non-selected populations

If the pair of selected populations diverged during the sweep phase, we modify our above predictions so that the lineages are in the same population for all of neutral phase I by substituting any term *t_I_* − *d_ij_* with *t_b_* (these terms represent the time during neutral phase I in which the pair of lineages are segregating in the same population). If the pair of selected populations are either the same population or diverged from each other during neutral phase II (when the selected allele segregates at frequency *x_s_*), and if the pair of lineages are linked to the same allele type at the most recent time they share a common ancestor, we consider the additional possibility that they coalesce during neutral phase II because of their associations with a subpopulation of haplotypes. We account for this using the same Poisson processes described above, except during neutral phase II the frequency of the selected background is xs and the frequency of the non-selected background is 1 − *x_s_*. Recall that in cases of earlier selection, it is possible that the common ancestor of a set of selected populations also has descendant populations that do not carry the allele at high frequency. For those populations, our models assume the selected allele has the same frequency trajectory as other selected populations until the time it diverges from them, after which we switch the frequency of the selected allele in this population to be its sampled frequency. This influences the probability that this population’s ancestral lineages were linked to the selected allele at the time its sweep phase begins.

In non-selected Neanderthal admixed populations, we track whether a lineage recombines out of its initial ancestry association, specified by its partition, over the entire time to introgression. If it loses this association, the only way it can have a non-neutral probability of coalescing with another lineage from a similarly non-selected population would be if that other lineage never recombined out of its initial background. As for a lineage from a selected population, they would have a non-neutral probability of coalescing if the lineage from the selected population maintained an association with a selected or non-selected background by the time of introgression. In the non-selected population, the probability that a lineage could have descended from Neanderthals, conditional on it recombining out of its background at sampling, is *g* throughout the whole time to introgression. We show how our predictions change with *t_b_* and *x_s_* in Figure S12 and Figure S13.

#### S3.3 Modifications if a population was sampled in the past

If a population was sampled in the past, we modify the probabilities that its sampled lineages are linked to each background according to the number of generations allotted to a given phase. For example, consider a population *i* that was sampled *τ_i_* generations ago, and carries the selected allele at high frequency such that in our application we consider it selected. If its sampling time falls within neutral phase II, then to calculate the probability that a lineage sampled from partition *B* of population *i* is linked to the selected background at the transition between neutral phase II and the sweep completion we simply reduce the amount of time for recombination in neutral phase II by *τ_i_*. If the provided waiting time until selection *t_b_* and selection coefficient s make it so that population *i* was instead sampled during the sweep phase, then at the transition between neutral phase II and the sweep completion, the sampled lineages are linked to the background denoted by their partition with probability 1. Then, we calculate that lineage’s probabilities of remaining associated with either background during the sweep phase as if the selected allele reached the lower frequency expected under the provided s and the allotted time for the sweep, *t_I_* − *t_b_* − *τ_i_*. If population *i* was sampled during neutral phase I, then we do not consider it a selected population, and it has *t_I_* − *τ_i_* generations for possible recombination by the introgression time.

#### S3.4 Incorporating migration among Neanderthal admixed human populations

In our application, most admixture graphs we use make CEU (the European ancestry representative population) a mixture of ancient populations WHG, EF, and Steppe. In our models, we modify predictions for partition *B* of CEU, assuming that ancient populations contributed only haplotypes with selected alleles to CEU. We thus consider donor populations to be only those contributing ancient populations with a partition *B* in the dataset. We set a single migration time *t_m_* to be the average migration time from all human donor populations (see Table S2). With probability exp(−*rt_m_*) a lineage sampled from partition *B* of CEU does not recombine before the migration time. In this case, the probability this lineage coalesces with a lineage sampled from any other partition *p_j_* is an average of the probability that each donor population’s partition *A* coalesces with the lineage from pj, weighted by the relative admixture proportions of each of these donor populations. If the lineage does recombine by the time of migration, the probability it coalesces with a lineage from pj is that of partition *b* of CEU. To predict coalescent probabilities for a pair of alleles sampled within CEU’s partition B, we average over results from the possible migration histories of both lineages. We acknowledge that our approach to treat migration does not reflect a realistic coalescent history. Other options did not allow our predictions to converge to CEU’s neutral probabilities of coalescing with increasing genetic distance from the selected site.

#### S3.5 Predictions for genotype partition *Bb*

After making modifications for migration, we make predictions for all Bb partitions. The probability a lineage sampled from this partition in population *i* coalesces with any other partition from any other population *p_j_* is the mean of this prediction for population i’s partition *B* and partition *b*. This reflects that half of the time we are sampling a neutral allele that is linked to the selected allele, whereas in the remainder of cases the neutral allele is linked to the non-selected allele. The probability of coalescing between partition *Bb* of population *i* and partition *Bb* of population *j* is the mean of the probability of coalescing of all four combinations of partitions A and a in populations *i* and *j*, in which there is a 50% chance either partition is sampled in either population. For probabilities of coalescing between lineages that were both sampled from partition Bb of population i, we take a weighted average of the probabilities of coalescing within partition *B*, within partition *b*, and between partition *B* and partition *b*, all partitions corresponding to population i. The first two cases occur with probability 0.25 and the last case occurs with probability 0.5.

**Figure S12:**
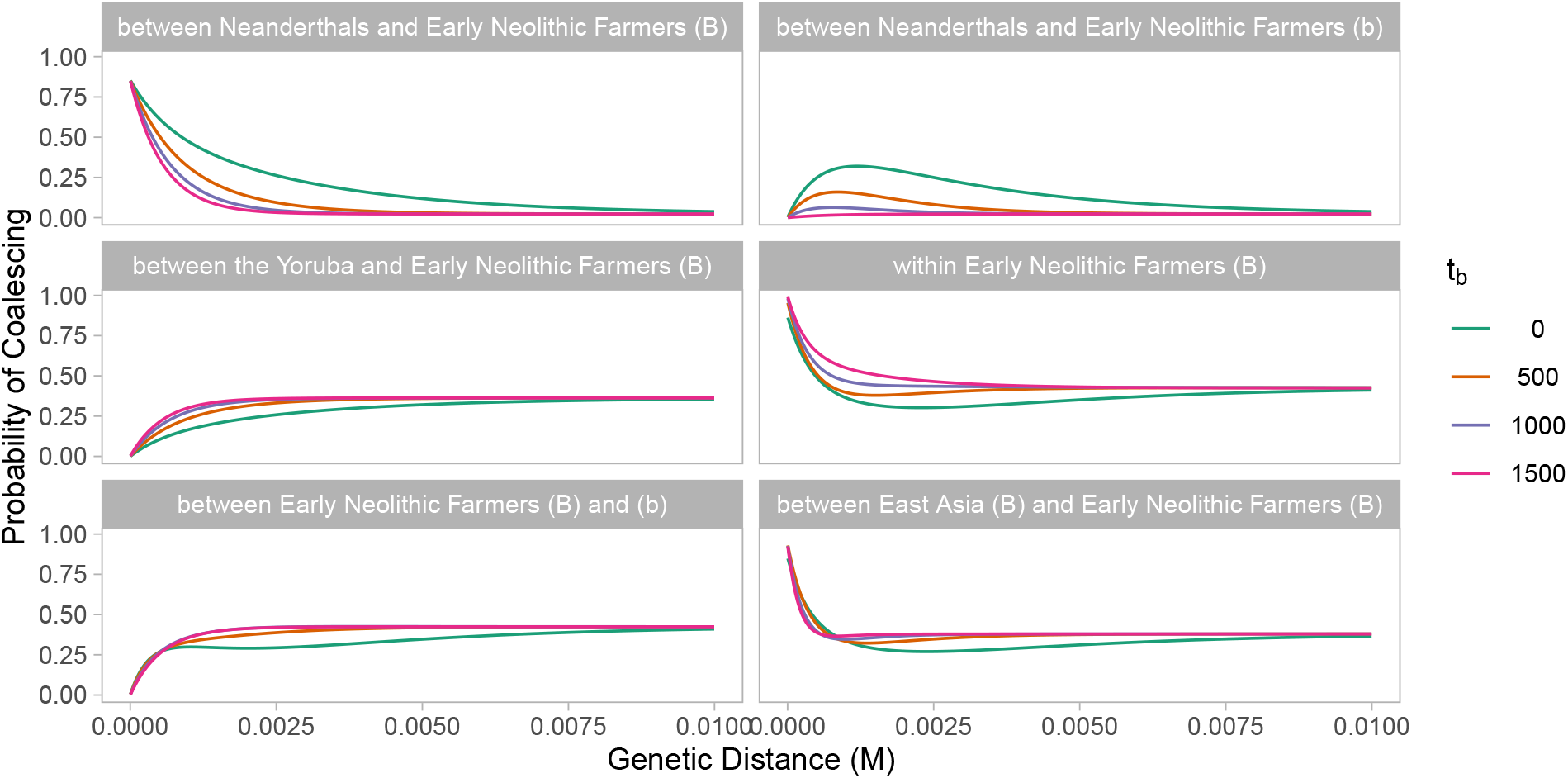
Model predictions under different values of (the waiting time until selection) when *s* = 0.01 and *x_s_* = 0.7.

**Figure S13:**
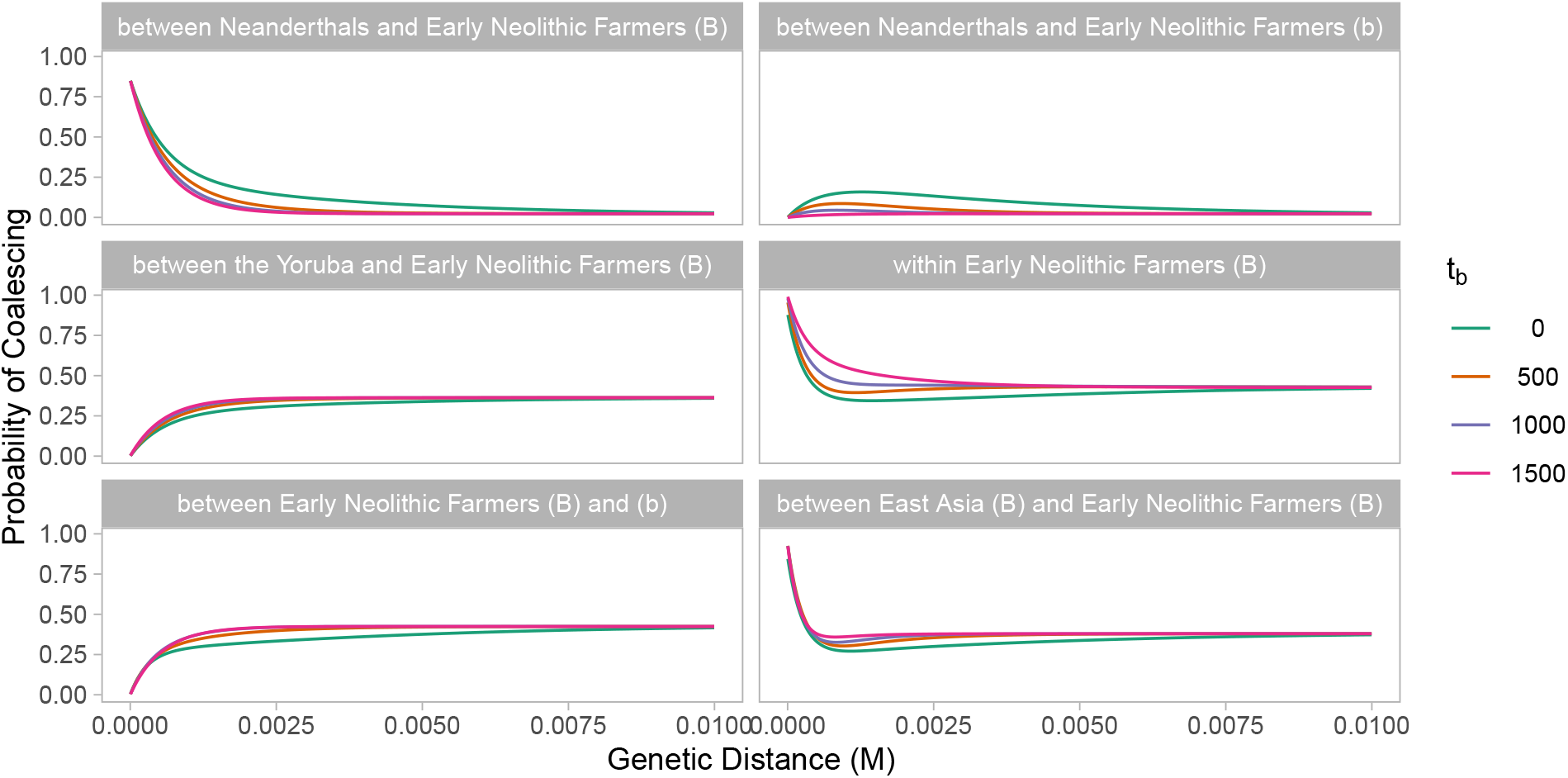
Model predictions under different values of (the waiting time until selection) when s = 0.01 and *x_s_* = 0.3. Note that predictions of become less distinguishable when *x_s_* decreases (see Figure S12 for comparison to *x_s_* = 0.7).

